# Tensor Image Registration Library: Automated Deformable Registration of Stand-Alone Histology Images to Whole-Brain Post-Mortem MRI Data

**DOI:** 10.1101/2022.08.11.503605

**Authors:** Istvan N. Huszar, Menuka Pallebage-Gamarallage, Sarah Bangerter-Christensen, Hannah Brooks, Sean Fitzgibbon, Sean Foxley, Marlies Hiemstra, Amy F.D. Howard, Saad Jbabdi, Daniel Z. L. Kor, Anna Leonte, Jeroen Mollink, Adele Smart, Benjamin C. Tendler, Martin R. Turner, Olaf Ansorge, Karla L. Miller, Mark Jenkinson

## Abstract

**Background:** Accurate registration between microscopy and MRI data is necessary for validating imaging biomarkers against neuropathology, and to disentangle complex signal dependencies in microstructural MRI. Existing registration methods often rely on serial histological sampling or significant manual input, providing limited scope to work with a large number of stand-alone histology sections. Here we present a customisable pipeline to automate the registration of stand-alone histology sections to whole-brain MRI data.

**Methods:** Our pipeline registers stained histology sections to whole-brain post-mortem MRI in 4 stages, with the help of two photographic intermediaries: a block face image (to undistort histology sections) and coronal brain slice photographs (to insert them into MRI space). Each registration stage is implemented as a configurable stand-alone Python script using our novel platform, Tensor Image Registration Library (TIRL), which provides flexibility for wider adaptation. We report our experience of registering 87 PLP-stained histology sections from 14 subjects and perform various experiments to assess the accuracy and robustness of each stage of the pipeline.

**Results:** All 87 histology sections were successfully registered to MRI. Histology-to-block registration (Stage 1) achieved 0.2-0.4 mm accuracy, better than commonly used existing methods. Block-to-slice matching (Stage 2) showed great robustness in automatically identifying and inserting small tissue blocks into whole brain slices with 0.2 mm accuracy. Simulations demonstrated sub-voxel level accuracy (0.13 mm) of the slice-to-volume registration (Stage 3) algorithm, which was observed in over 200 actual brain slice registrations, compensating 3D slice deformations up to 6.5 mm. Stage 4 combined the previous stages and generated refined pixelwise aligned multi-modal histology-MRI stacks.

**Conclusions:** Our open-source pipeline provides robust automation tools for registering stand-alone histology sections to MRI data with sub-voxel level precision, and the underlying framework makes it readily adaptable to a diverse range of microscopy-MRI studies.

**Highlights:**

- New software framework for prototyping bespoke image registration pipelines
- Automated pipeline to register stand-alone histology sections to whole-brain MRI
- Novel deformable slice-to-volume registration algorithm
- No strict necessity for serial histological sectioning for MRI-histology registration

## 1. Introduction

MRI is a powerful neuroimaging technique providing non-invasive images of the entire brain but suffers from limited spatial resolution and biological non-specificity. In comparison, microscopy techniques are highly complementary, conferring specificity through high spatial resolution and precise targeting of cellular constituents, but being highly invasive (e.g., requiring tissue extraction). Combined MRI-microscopy studies are useful for validating radiological signs of disease against neuropathological evidence [1, 2], and to improve biophysical models [3] that infer microstructural properties of the tissue beyond ordinary resolution limits of MRI.

Depending on the aim, MRI-microscopy datasets can vary along many axes: 1) whole brain [4] vs tissue blocks [5], 2) serial histological sectioning [6] vs single-section sampling [7], 3) large [8] vs small [9] histology sections, 4) ex-vivo [10] vs post-mortem MRI [11], and 5) the exact combination of MRI and microscopy modalities used. This diversity of the input data presents unique challenges [12] for the alignment of MRI-microscopy images (e.g., extreme contrast differences, vastly different spatial resolutions, 2D vs 3D image domains), which are difficult to overcome with existing registration software that was not optimised for this task. This is especially true if the source code is closed, or an inflexible implementation prohibits the customisation of the core algorithm.

An overwhelming majority of previous works addressed microscopy-to-MRI registration via volumetric reconstruction of serial sections [13–31]. A comprehensive review of these techniques was published by Pichat et al [32]. For these methods, the tissue must be sectioned with a constant slice gap. First, the histology images are undistorted in 2D using photographs of the tissue block as a reference. Subsequently, the undistorted histology images are stacked to create a volume, which is then registered to MRI using 3D registration tools such as ABA [33] or ANTs [34]. While the results are highly accurate, these methods cannot work with single-section histology images, and serial histological sampling is often prohibitively labour-intensive, especially for whole-brain coverage in multiple subjects [35].

Registering stand-alone histology images to volumetric MRI data on the other hand presents unique challenges. First, the 2D-to-3D transformation must account for both the in-plane deformations of the tissue section as well as the bulk deformations of the brain that may deflect the sectioning plane. Second, a complex transformation model implies a vast parameter space, that must be navigated effectively during the optimisation to find the global optimum. Third, the cost function must be able to account for the contrast, data type, and dimensionality difference of the input images. Finally, the algorithm should aim to be fully automated, (e.g., without requiring manual landmarks) to be feasible for larger datasets. A comprehensive survey of slice-to-volume registration methods was published by Ferrante et al [36]. One early landmark-free approach used 2^nd^ and 3^rd^-degree polynomial extensions of the 3D affine transformation model [37] but achieved limited accuracy (3-8 mm) [38]. Meyer et al [7] introduced a thin-plate spline (TPS) transformation model and obtained “visually accurate” results. A comprehensive study by Osechinskiy et al [39, 40] concluded that transformation models, cost functions, and optimisation methods must be tailored to the specifics of the input MRI and histology data. The additional challenge of registering small-format histology sections (as opposed to whole-hemisphere sections) was later addressed by Ohnishi et al [41], using manual landmarks to stitch together multiple histology images and register them indirectly to MRI via a brain slice photograph. Goubran et al [42] introduced a hybrid 2D/3D algorithm specifically for sparsely sampled histology sections that alternates between slice-based and volume-based registration with ex-vivo MRI. However, their method relies on multiple slices and cannot account for 3D slice deformations. While these works collectively laid down important algorithmic foundations, each of them concerned a specific problem at hand, and the underlying software framework was not released to the wider community for further testing and refinement. HistoloZee [43] is a recent development that addresses the previously unmet need for histology-to-MRI registration software, and even provides an interactive graphical user interface. However, the registration process strongly relies on manual input, the transformation model cannot account for deformations of the sectioning plane, and the source code is closed.

Hence, existing software tools are not well-positioned to automate the registration of sparsely sampled histology sections to volumetric MRI data. An experimental MRI-microscopy registration framework is needed, that is open-source, and provides enough flexibility to create, test, and refine various algorithms. Simultaneously, the framework should exhibit a sufficiently high-level programming interface such that bespoke MRI-microscopy pipelines can be deployed in a timely manner. Ideally, one would additionally reduce the steep learning curve that is normally associated with the more general-purpose, low-level frameworks, such as the Insight Toolkit [44].

In this paper we describe a novel pipeline for the registration of sparsely sampled single-section histology images to MRI volumes of the human brain. A significant proportion of the pipeline is automated, and it is implemented in our newly built software framework, the Tensor Image Registration Library (TIRL). TIRL aims to provide a flexible solution for implementing bespoke image registration pipelines for diverse MRI-microscopy applications.

## 2. Methods

### 2.1. Registration workflow in the Tensor Image Registration Library

The Tensor Image Registration Library (TIRL) is a an open-source software platform (https://git.fmrib.ox.ac.uk/ihuszar/tirl, also distributed with FSL v6.0.4 and above) for implementing bespoke image registration routines in Python 3. We designed it for situations where the type of the input data (e.g., histology formats) or the nature of the registration problem (e.g., 2D-to-3D transformation) makes it difficult to employ existing software, and one can benefit from taking full control over the registration process with the granularity of individual parameter updates. TIRL is highly modular; it consists of generic objects that may be customised (via parameters or subclassing) and assembled in unique ways within a Python script – hereafter designated as a *TIRL script* – to perform specialised image registration tasks. While we summarise the main design concepts of TIRL here, readers may refer to a full documentation of the library at https://git.fmrib.ox.ac.uk/ihuszar/tirldocs.

Figure 1 shows the anatomy of a basic TIRL script that registers two images. A core part of the library is a universal image container, the TImage object (Figure 1, *black box*). Image data is imported from disk in chunks into the TImage to avoid memory overload. Image data in the TImage is defined on an *N*-dimensional discrete manifold (a grid or scattered datapoints), where each datapoint can be an *L*-rank tensor (scalar/vector/matrix/tensor). For interim points, image data is retrieved by the associated Interpolator object (Figure 1, *yellow box*), supporting nearest-neighbour, linear, and spline interpolation by default. (For large images, interpolation is internally distributed across parallel processes for higher performance.) Each TImage has an associated Domain (Figure 1, *red box*), which represents the pixel/voxel coordinates of the image. Pixel/voxel coordinates are mapped to physical coordinates by the Chain of Transformation objects that is assigned to the Domain. The Chain is divided into two parts. The *internal* Chain (Figure 1, *grey parallelograms*) is managed by TIRL to store the resolution of the image and to preserve the physical coordinates of the image when padding is applied. The *external* Chain is where the author of the TIRL script can specify an arbitrary sequence of linear and non-linear Transformation objects (Figure 1, *white boxes*) for optimisation. The parameters of the external Chain can be optimised either all-at-once or in arbitrary groups, identified by OptimisationGroups (Figure 1, *brown box*). The registration process is controlled by the Optimiser object (Figure 1, *green box*), which iteratively updates the selected parameters within their predefined range according to its own predefined algorithm. The Optimiser’s objective function is evaluated at every iteration as a sum of image-specific cost and parameter-specific regularisation terms, represented by the respective Cost (Figure 1, *blue box*) and Regularisation objects (Figure 1, *light grey box*).

**Figure 1.**
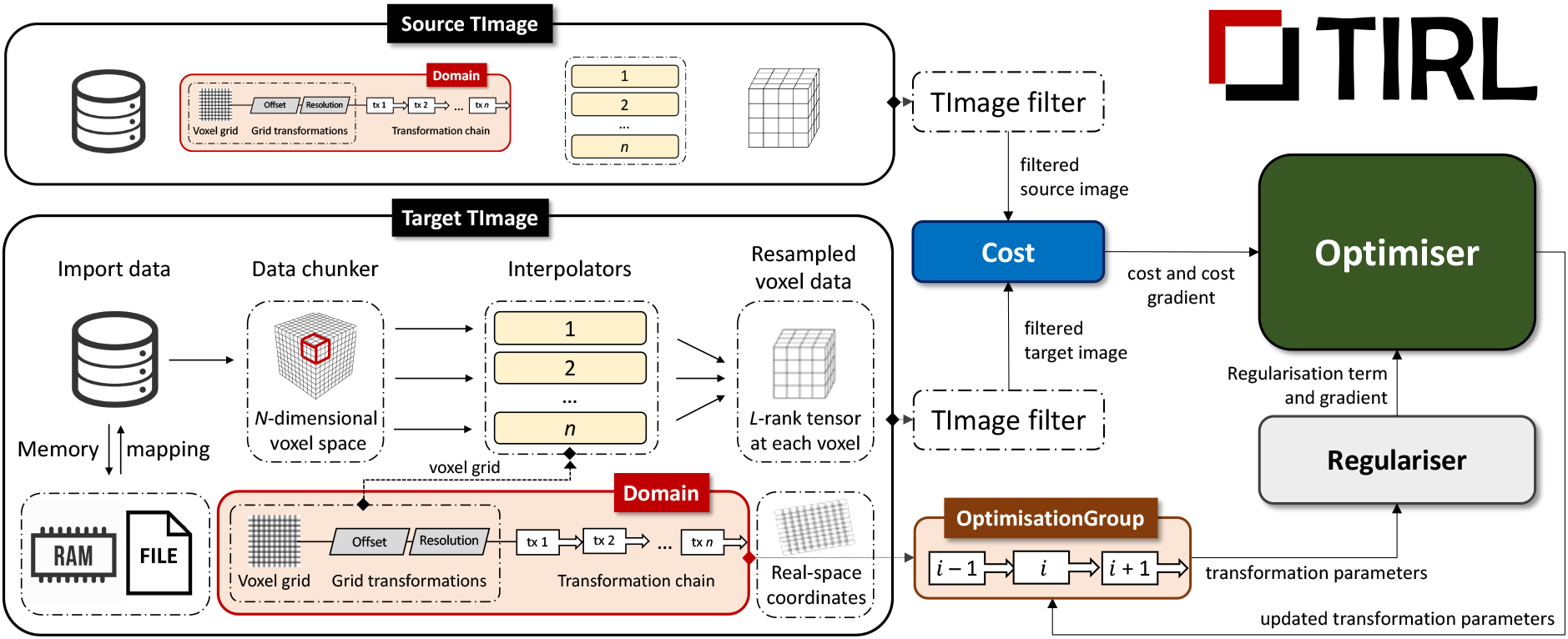
Schematic representation of a generic TIRL registration workflow. Specialised instances of this workflow are implemented by all four stages of the pipeline, employing specialised subclasses of the Cost, Optimiser and Regulariser base classes. For a detailed description of the objects/classes, the reader is referred to the general documentation of TIRL. A coarse overview of the TImage object and the workflow is given in section 2.1 of the main text.

Using the above scheme, one can create individual TIRL scripts that specialise in a specific type of input or transformation, then assemble these into a bespoke modular registration pipeline by passing the optimised transformation chain from one script to another. Any object of the workflow can be saved into a TIRL file or loaded from a TIRL file at any time, which eliminates compatibility issues, and makes it straightforward to interrogate the results even at the level of elementary transformations. TIRL transformation chains can be split and freely recombined, as well as concatenated with FLIRT [45, 46] matrices or FNIRT [47] fields, providing full interoperability with FSL [48] registration tools. Finally, TIRL chains have built-in methods to realign vectors and tensors under transformations, making them compatible with direction-sensitive data, such as diffusion MRI.

In the following sections, we overview the TIRL scripts that we created to register histology sections to MRI data in an existing dataset. We implemented these in a general style, with several configuration options, with the aim of making them directly accessible to users without professional coding skills. The scripts are distributed as part of a growing open-source collection, called TIRLScripts (https://git.fmrib.ox.ac.uk/ihuszar/tirlscripts).

### 2.2. MRI-histology dataset

For demonstrating histology-to-MRI registration with TIRL, we resourced images from a previous post-mortem study [49], only including subjects with a consistent set of histology, photographic, and MRI data (Figure 2). All data was collected and used according to the Oxford Brain Bank’s (OBB) generic Research Ethics Committee approval (15/SC/0639). Written informed consent was obtained by the OBB from all participants of this study. The image acquisition [49–51] and post-processing [52–54] details have been described earlier; we only summarise the most important aspects here.

**Figure 2.**
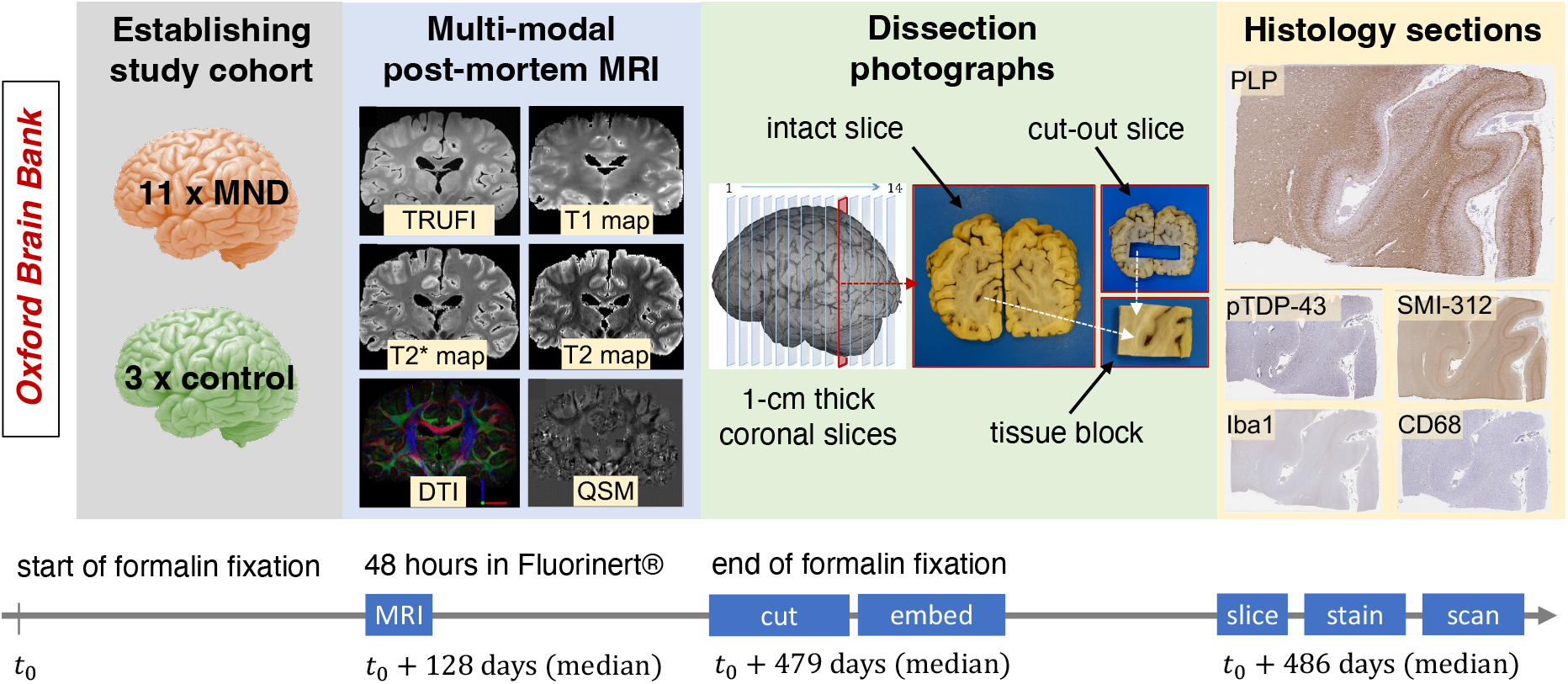
Overview of the MRI-histology dataset for demonstrating histology-to-MRI registration with TIRL. Fourteen (11 MND + 3 control) post-mortem brains with a consistent set of multi-modal MRI data, dissection photographs, and digitised histology slides were resourced from a previous study [49]. Further details are given in section 2.2 of the main text.

Our reduced dataset represented a mixed group of post-mortem brains, 11 of which were affected by terminal-stage motor neuron disease (MND) and 3 brains without pathological evidence of neurodegeneration at the time of death (median age: 65.5 years). The brains were immersed in 10% neutral buffered formalin for a median duration of 4 months.

MRI scans (Figure 2, blue panel) were acquired on a 7T Siemens Magnetom scanner and processed to produce quantitative T1 and T2 maps at 1 mm isotropic resolution, T2* and susceptibility maps at 0.5 mm isotropic resolution, diffusion-derived parametric maps at 0.85 mm isotropic resolution, and a TRUFI anatomical reference scan at 0.25 mm isotropic resolution. All modalities were aligned to TRUFI space using FSL’s Linear Registration Tool (FLIRT), and to the 1 mm MNI152 template using ANTs.

As further shown in Figure 2 (green panel), the brains were subsequently dissected by hand to create approximately 1 cm thick coronal sections, starting from the plane of the mammillary bodies. The total number of slices (13-17) varied with the size of the brain. One or more (usually 4-8), approximately 25 × 35 mm large tissue blocks were sampled from predefined anatomical locations of each coronal section. The tissue block sampling process was carefully documented by taking photographs of both sides of the coronal slices and the extracted tissue blocks. The brain slices were photographed repeatedly, whenever a new block was sampled from them to create a series of “cut-out” images. Photographs were 5472 × 3648 pixels large (approximately 50 μm/pixel).

The tissue blocks were embedded in paraffin and sectioned on their anterior surface on a microtome at 6-10 μm thickness. Consecutive tissue sections from each block were immuno-stained separately for myelin proteolipid protein (PLP), neurofilaments (SMI-312), microglia (Iba-1), activated microglia and macrophages (CD68), and phosphorylated TAR-DNA binding protein-43 (pTDP-43), and counter-stained with haematoxylin to visualise cell nuclei [49]. The slides were digitised in SVS format using an Aperio ScanScope slide scanner at × 20 objective magnification, yielding a typical image size of 60,000 × 45,000 at full resolution (approximately 0.5 μm/pixel) and thumbnails at 8 μm/pixel resolution (Figure 2, yellow panel).

### 2.3. Creating a multi-stage TIRL pipeline for histology-to-MRI registration

Histology sections are prone to distortions, and it is often very difficult to localise them in whole-brain MRI data without anatomical knowledge. We eliminate these difficulties and automate most of the registration process by proposing a multi-stage pipeline (Figure 3), that relies on two intermediate photographs to undistort (Stage 1) and guide the insertion (Stages 2 & 3) of each histology image into MRI space.

**Figure 3.**
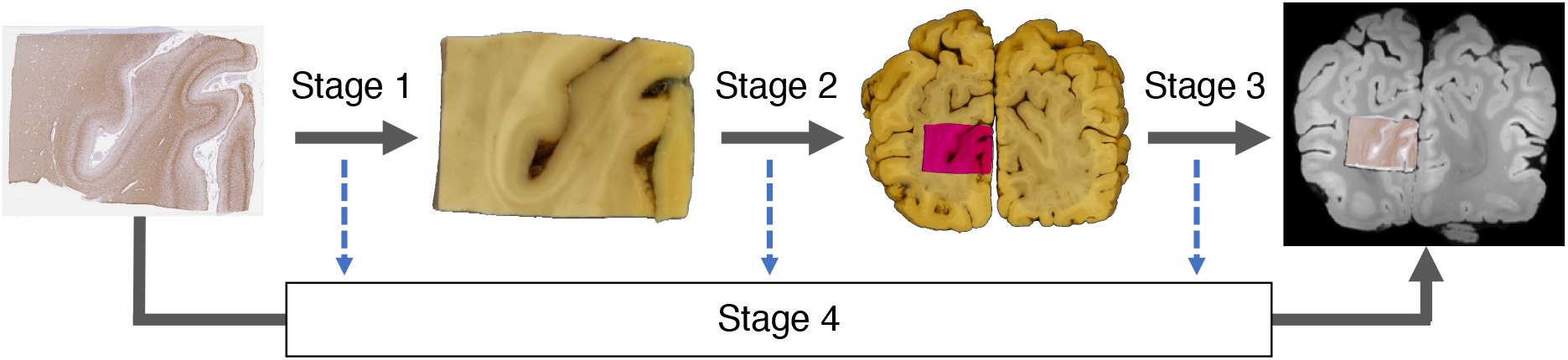
Overview of the automated histology-to-MRI registration pipeline with two photographic intermediaries. Each stage maps the pixel coordinates of the input image to the pixel/voxel coordinates of the output image by a chain of transformations. The stage-specific transformation chains are optimised separately and eventually combined to obtain a one-to-one (invertible) mapping between histology and MRI. Due to the generality of the transformations, each histology image is mapped onto a *parametric surface* in MRI space. Images are not shown to scale.

Figure 3 shows how the histology-to-MRI transformation chain is broken down into three independent parts, whose optimisation may be carried out in parallel (Stages 1-3). The optimised stage-specific chains are then concatenated and fed into a 4^th^ stage to refine the transformation parameters by directly registering histology to MRI data. Each stage is implemented as a stand-alone Python script, which uses the components and methods of the TIRL package and is accompanied by a YAML-formatted configuration file, allowing users to customise each stage for their own data. For further details, readers can refer to the openly available source code of the scripts and annotated configuration files (https://git.fmrib.ox.ac.uk/ihuszar/tirlscripts). Example data from the above-mentioned MRI-histology dataset with completed registrations are available through the Digital Brain Bank (https://open.win.ox.ac.uk/DigitalBrainBank/#/datasets/pathologist). The full dataset is available on request via a material transfer agreement to ensure that the data is used for purposes that satisfy research ethics and funding requirements.

In the subsequent sections, we discuss the multi-stage optimisation sequence of the transformation chain that maps the pixels of a stand-alone histology onto the voxels of whole-brain MRI, and further discuss the experiments to validate the accuracy of each stage.

### 2.4. Stage 1

The goal of Stage 1 is to establish a forward mapping from the pixel coordinates of a histology image to the pixel coordinates of the corresponding tissue block photograph (Figure 3). Stage 1 therefore accounts for the deformations of the tissue section that occur while it is mounted on the glass slide. Both images are pre-processed before the registration in line with the Stage-1 configurations.

#### Pre-processing

The block photo was cropped loosely around the edges of the block (to eliminate other objects from the frame) and background-segmented by pixelwise *k*-means clustering (*k*=2) using auxiliary scripts (provided via Git). The Stage-1 script automatically downsamples the histology image by a Gaussian kernel (FWHM = 6.25 pixels) to equalise the resolution of the inputs. Both images are converted to grayscale. The histology image is padded on all four edges by 1/6^th^ of the respective image dimension. Padding avoids trivial reductions in cost function by simply shifting one image outside the other’s field of view. A manually defined mask is occasionally supplied as an input with the histology image to exclude artefactual drivers of the registration, such as tears, holes, folds, stain deficiencies, overstaining, tissue debris, bubbles, or slide scanning defects. To bridge the modality gap, equal representations of the images are obtained by applying a non-linear filter, the Modality Independent Neighbourhood Descriptor (MIND) [55] to the grayscale images. MIND accentuates edges in the images by replacing pixel values with an 8 × 1 vector describing the intensity relationship of the pixel with its immediate neighbours. The images are initially aligned by their geometrical centres.

#### Registration

The Stage-1 chain consists of the following transformations (Figure 4, chain): 2D rotation (about the geometrical centre of the histology image), isotropic scaling, 2D translation, 2D affine, and a pixelwise defined displacement field. The registration cost (hereafter referred to as *MIND cost*) is calculated as the sum of pixelwise Euclidean distances of MIND vectors across the histology image domain. The MIND cost is successively minimised in 3 linear and 1 non-linear step (Figure 4). The linear registration steps uniformly employ the gradient-free bounded BOBYQA optimisation method [56]. For the non-linear registration, the cost function is extended with a diffusion regularisation term [55, 57] that enforces smooth deformations by penalising sharp gradients in the displacement field. The relative weight (a) of the regularisation term is determined empirically for each dataset. The displacement vectors are initialised to 0 and refined by 20 or fewer iterations of Gauss–Newton optimisation [55, 58] at each of the prespecified resolution levels (typically 0.8, 0.4, and 0.2 mm/pixel).

**Figure 4.**
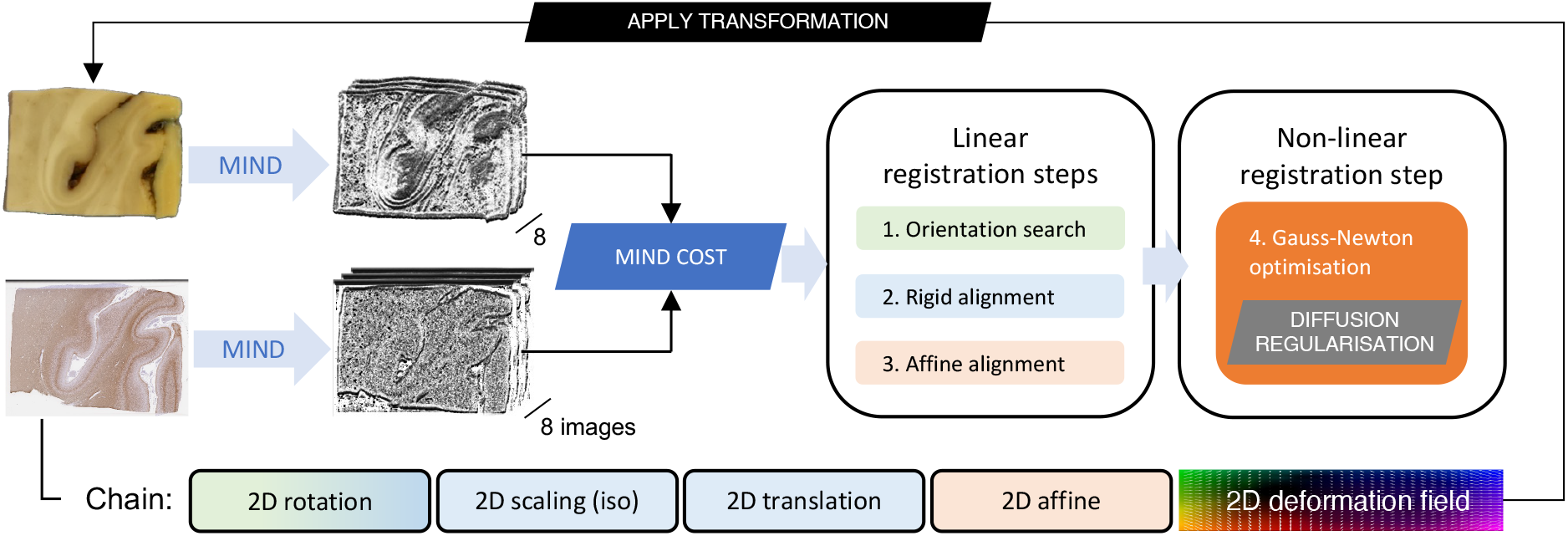
Stage 1 – deformable registration of a histology image to a tissue block photograph. Contrast differences between the input images are equalised by applying the non-linear image filter MIND. Image dissimilarity is defined as the Euclidean distance between the MIND representation of the images. The parameters of the Stage-1 transformation chain are found in three successive linear and one non-linear optimisation steps. See further details in section 2.4 of the main text.

#### Experiments

We performed Stage-1 registrations on PLP-stained histology images from the hippocampus and anterior cingulate cortex regions of all subjects (2 × 14 images). A foreground mask was generated for the block photos by thresholding at 0.1 relative intensity and dilating by a 10×10 pixel kernel. The regularisation weight was empirically set to a = 0.4.

To quantitatively assess the accuracy and robustness of the registration, ground-truth grey-white matter contours were segmented by hand on both the original tissue block photographs and the histology images. The histology contours were transformed by the optimised Stage-1 chain and compared with the respective photographic contours by calculating the median contour distance (MCD, in millimetres). MCDs were compared between the linear and non-linear registration steps and plotted for different regularisation weights. Finally, the accuracy of the Stage-1 registration was compared for both anatomical regions against various ANTs paradigms, including both the Mattes mutual information and the cross-correlation metrics that were used in a previous study [25] to register histology sections. Further details of the ANTs registration parameters are given in Supplementary material 1.

### 2.5. Stage 2

The goal of Stage 2 is to establish a forward mapping from the tissue block photograph to the corresponding coronal brain slice photograph (Figure 3). This stage eliminates the need for anatomical knowledge to manually localise small tissue sections within whole-brain MRI data. Both images are pre-processed before the registration in line with the Stage-2 configurations.

#### Pre-processing

The pre-processing steps for the input images are identical to those in Stage 1 with respect to cropping, background segmentation. The Stage-2 script also converts inputs to grayscale.

#### Sampling site determination

All photographs pertaining to a specific coronal slice of the brain are collected and automatically sorted starting from the most intact image of the slice towards the slice with the most regions missing. Consecutive image pairs (Figure 5G-K) are aligned by a succession of rigid, affine, and non-linear registration, as described in Stage 1. The non-linear registration is carried out at slightly coarser resolutions (1.5 mm/pixel and 1 mm/pixel) and with higher regularisation (α = 0.6) to avoid excessive deformations around missing regions, but no masks are used at this stage. The aligned image pairs are binarized and their difference (XOR) is taken to highlight potential sampling sites (Figure 5M). Sites with area smaller than 1 cm^2^ or width narrower than 4 mm are considered minor registration errors, and hence discarded. The centroids of the remaining blobs are deemed possible sampling sites, and their coordinates are mapped onto the most intact slice (Figure 5G), which are then used as the registration target for the individual blocks (e.g., the one in Figure 5L).

**Figure 5.**
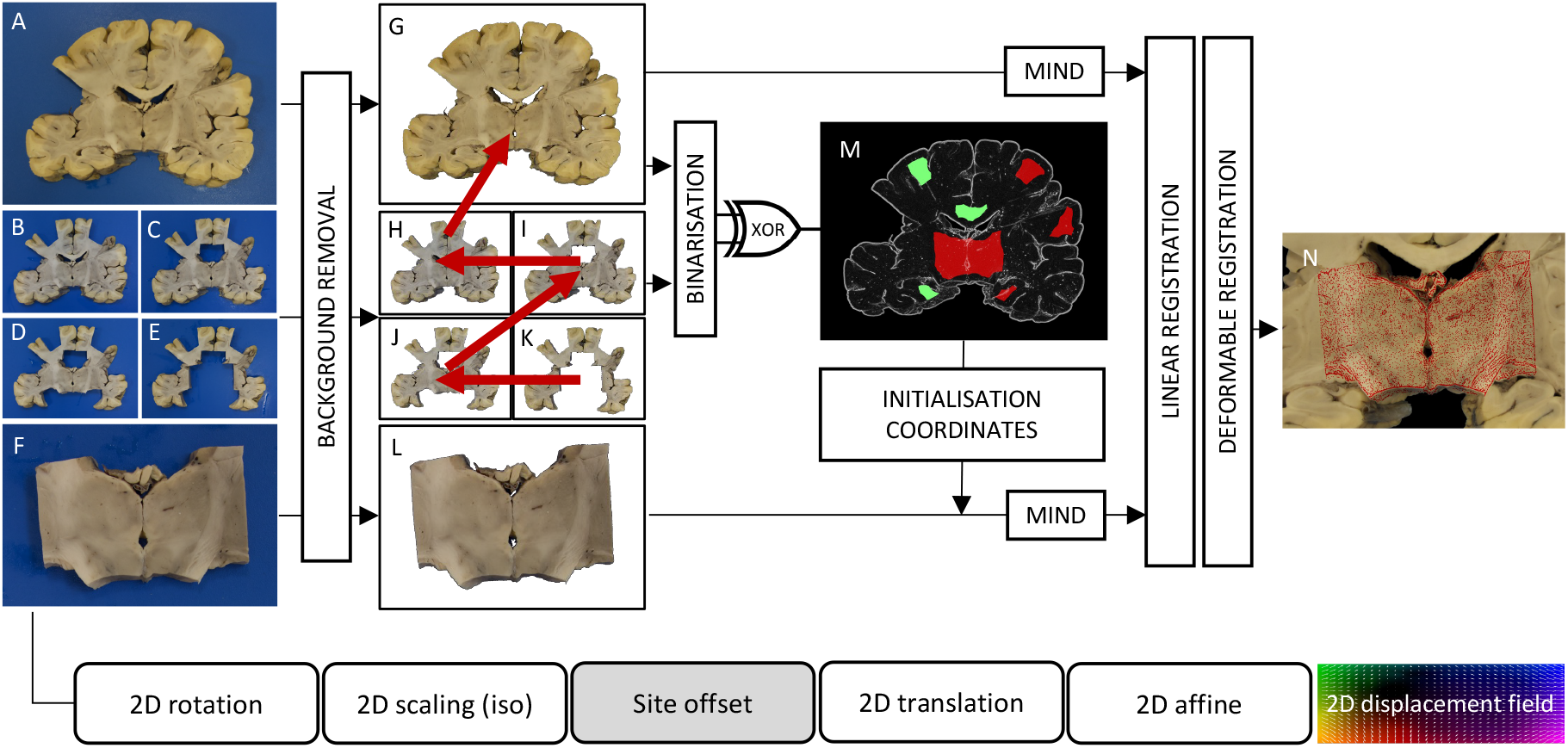
Stage 2 – Automated sampling site matching and deformable registration of tissue blocks to coronal brain slices. The raw inputs (A-F) are background-extracted (G-L), and the sampling sites on G are automatically identified by binarizing and pairwise subtracting (XOR) subsequent photographs of the coronal brain slice (G-K). Tissue block photographs (F) are cross-matched with the identified sampling sites (M) and their alignment is fine-tuned (N) at the relevant site using both linear and diffusion-regularised deformable registration.

#### Registration

The Stage-2 chain maps the pixel coordinates of the tissue block onto the pixels of the corresponding brain slice photograph. The Stage-2 chain consists of the following transformations: a 2D rotation (about the centre of the tissue block), a 2D isotropic scaling, a fixed 2D translation to the sampling site (sampling site offset), a variable 2D translation, a 2D affine, and a pixelwise displacement field defined over the domain of the tissue block image (Figure 5). To identify the correct sampling site for any block, the block is first initialised at all sites, and the MIND cost is evaluated for set combinations of rotations (typically in 30-degree increments) and translation parameters (typically at +/−10 mm from the sampling site) to account for small inaccuracies of the sampling site determination. The best three sets of parameters at each site are fine-tuned by BOBYQA optimisation, and the one associated with the lowest MIND cost at the end of this process is used to initialise the chain by setting the sampling site offset and the 2D rotation. From this initial state, the registration proceeds through rigid, affine, and non-linear optimisation as described in Stage 1 to fine-tune the rest of the Stage-2 chain parameters. Masks to exclude the background are used throughout all Stage-2 registration steps and are generated automatically by thresholding both grayscale inputs at 10% relative intensity.

#### Experiments

We performed Stage-2 registrations on 87 tissue blocks from various anatomical regions (corpus callosum, anterior cingulate cortex, hippocampus, visual cortex). To test the accuracy of the registration, contours were defined manually along salient anatomical features on 28 image pairs and the MCD were measured after registration. The robustness of the automatic sampling site matching was tested by registering 8 blocks that were extracted from the same brain slice. Finally, to test the robustness of Stage-2 registration against block initialisation error, we simulated the registration of the same 8 blocks from 100 different positions around the centre of their respective sampling sites and counted successful registrations (<0.2 mm MCD) as a function of initialisation error in millimetres.

### 2.6. Stage 3

The goal of Stage 3 is to establish a forward mapping from the pixel coordinates of a coronal brain slice photograph to the voxel coordinates of an MRI volume (Figure 3). Crucially, we make very few assumptions about the physical brain slices in Stage 3. While their orientation is ‘coronal’, it is unlikely that they correspond perfectly to acquisition slices of the MRI data. In fact, it is possible that the sectioning plane is curved in MRI space, due to the irregularity of the cuts or the bulk deformations of the brain during either dissection or scanning. Therefore, Stage 3 leverages the unique cross-sectional anatomy of the brain slice photographs (e.g., the shape of the cortical ribbon, the cross section of subcortical nuclei and ventricles) to find a 3D surface in MRI space that best represents the “cutting plane” and maps the pixels of the 2D slice photograph onto this surface. Slight in-plane deformations of the brain slices are also taken into account, as they could jeopardise the alignment of anatomical structures. As the number of transformation parameters is large, Stage 3 makes extensive use of parallel computing by performing grid searches, ranking interim results, and employing nested gradient-free local optimisations, which make it the most algorithmically complex part of the entire pipeline. The Stage-3 algorithm was developed empirically in a detailed trial-and-error process. The schedule of parameter re-initialisations and optimisation bound updates was found to be critical to achieve general robustness. While this may give the impression that Stage 3 would be difficult to use with a different dataset, in practice we found the current implementation to be readily adaptable for a range of microscopy and MRI images of both mouse and macaque brains by changing the concomitant Stage-3 configuration file. Readers may compare the different configurations that are provided in the Git repository to learn more about adapting Stage 3. Here, we provide a high-level overview of the optimisation process, which is common to all protocols.

#### Pre-processing

The brain slice photograph was cropped and background-extracted before importing to Stage 3 using auxiliary scripts (provided via Git). The Stage-3 script converts the input to grayscale, and downsamples it by a Gaussian kernel (FWHM = 5 pixels) to match the resolution of the MRI (0.25 mm/voxel). Where necessary, slice masks were generated by hand to exclude areas where the brain slice had been damaged due to further investigations on the motor cortex (Supplementary material 2).

#### Registration

The Stage-3 chain consists of the following transformations (Figure 6): a 2D isotropic scaling, a 2D rotation about the slice centre, a 2D translation, a 2D-to-3D embedding (sets z = 0), a 3D displacement field, a 3D rotation (about the adjusted centre of the slice photograph), a 3D translation, and a 3D affine. The chain parameters are initialised such that the brain slice corresponds to the middle layer of a 2-cm-thick rectangular slab (Figure 6, panel 1, in orange). The slab, which is defined manually in the configurations by its centre and orientation, represents the spatial extent of the 4-step optimisation process. The first step (Figure 6, panel 1, *rigid search*) moves the centre of the photo to several (usually 5-11) equidistant points along the central axis of the slab and varies the 3D rotation parameters at each of these in a prespecified range (e.g., 3 values in a 30-degree range about each axis). Each combination of the initial rigid parameters is then refined in a multi-resolution local optimisation scheme (typically 2, 1, 0.5, and 0.25 mm/pixel) that minimises the MIND cost. The MIND cost is always calculated between the brain slice photo and the MRI data that is resampled onto the same 2D domain. This step employs heavy parallelisation and interim results are constantly ranked to reduce the number of optimisations that need to be carried out at higher resolutions. The second step (Figure 6, panel 2, affine alignment) starts from the best rigid position and orientation of the slice and optimises the 3D affine matrix to account for shears. The last two steps estimate the vectors of the 3D displacement field: first the in-plane components only (Figure 6, panel 3), then both the in-plane and the orthogonal components simultaneously (Figure 6, panel 4). As an empirical compromise between accuracy and computational efficiency, exact displacements are estimated for only a small number of evenly distributed control points (typically 32), which are generated automatically by the script. For the rest of the pixels, the local displacements are calculated by interpolation using Gaussian radial basis functions. All optimisations throughout Stage 3 employ the BOBYQA method and minimise the MIND cost, which demonstrated superior robustness in our experiments when compared to normalised mutual information (Supplementary Material 3).

**Figure 6.**
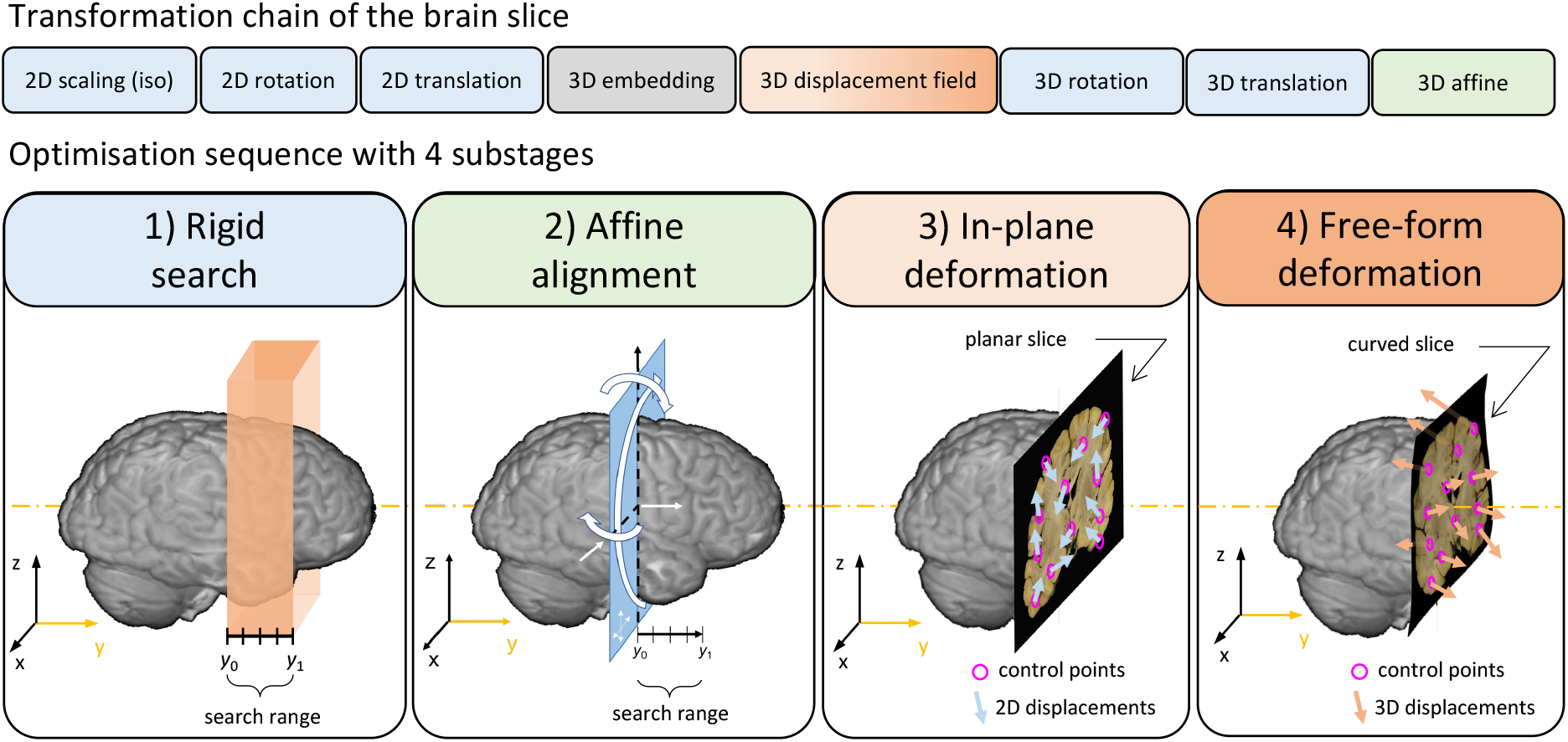
Stage 3 – Deformable registration of a brain slice photograph to an MRI volume. The four tiles from left to right illustrate consecutive steps of the optimisation process. See the main text for further details.

#### Experiments

To assess the accuracy and robustness of Stage 3, we registered 209 brain slice photographs from 14 subjects (approximately 15 slices per subject), and inspected the alignment of salient anatomical structures, with special attention to the highly variable grey-white matter boundary, ventricle cross sections, and perforating vessels.

#### Simulations

To quantify the accuracy of Stage 3, and specifically its ability to compensate 3D deformations of 2D slices, we also performed registrations with simulated slices. These were generated by virtually recreating the coronal slicing scheme (Figure 2, *green panel*), i.e., by resampling the structural MRI data of a single subject onto a series of analytically defined parallel first-order (planar) and second-order (quadratic) polynomial surfaces. Two groups of planar and quadratic slices were generated: first in coronal orientation, then slightly (10 degrees) tilted towards the left and inferior directions for increased difficulty. Starting from a perturbed position and orientation, the slices were registered to structural MRI data by the Stage-3 algorithm. We calculated the median registration error (MRE) for each slice and for each optimisation step by measuring the median distance of the registered slice pixels from the corresponding analytical surface points.

### 2.7. Stage 4

The goal of Stage 4 is to fine-tune the alignment of a histology image after it has been registered in MRI space by the previous three stages. Stage 4 accounts for two specific imperfections of the intermediate photographs, which are discussed below in conjunction with the most important methodological considerations for this stage.

#### Imperfection of the tissue block photographs

Anatomical inconsistencies may arise between the tissue block photograph and the histology section if the histology section comes from several hundred micrometres deep inside the tissue block (as a result of shaving off an excessive number of tissue layers in the microtome). Such inconsistencies may drive the non-linear registration in Stage 1 to overestimate local deformations, leading to the misalignment and an overly distorted appearance of the histology image in MRI space.

#### Imperfection of the coronal brain slice photographs

Excessive widening or closing of the interhemispheric fissure (Figure 7A-B) in the brain slice photographs (compared to their relative configuration in the MRI volume) requires the estimation of local large displacements in Stage 3. However, this goes against the design principles of Stage 3, which defines the 3D displacement field sparsely to prevent local large deformations under the assumption that they are physically implausible.

**Figure 7.**
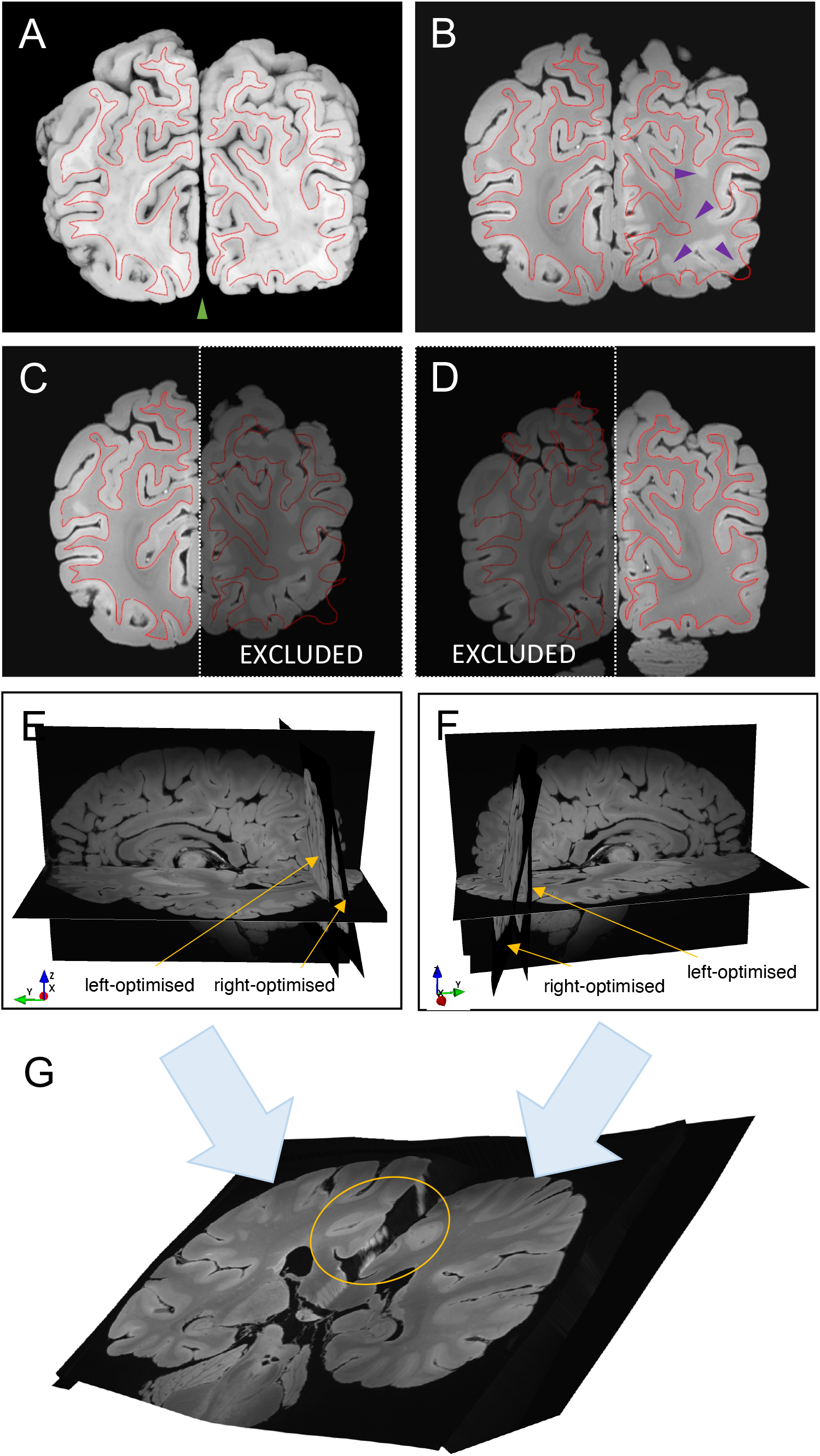
Stage 4 pre-processing – hemisphere-specific deformable registration of a coronal brain slice photograph to a structural MRI volume. The *red contour* represents the grey-white matter boundary as it is seen on the brain slice photograph **(A)**. Due to the closing of the interhemispheric fissure (*green arrowhead*), the bilaterally driven Stage-3 registration result **(B)** is not uniformly accurate (*purple arrowheads*). **(C-D)** Stage-3 registrations with hemisphere-specific masking produce accurate results. **(E-F)** Hemisphere-specific Stage-3 registration reveals large differences in the slicing plane between the left and the right hemispheres, which is most likely caused by the antero-posterior shearing of the hemispheres during dissection. **(G)** Merging hemisphere-specific slice-to-volume transformations results in a single smooth transformation of the slice that preserves the accuracy of the alignment in both hemispheres irrespective of variations in the interhemispheric gap or the antero-posterior shearing of the hemispheres (*encircled*).

#### Pre-processing

As a preparation for Stage 4, we run Stage 3 twice with hemisphere-specific 3D masks (Figure 7C-D) to maximise the registration accuracy within the hemispheres. Using an auxiliary script (provided via Git) we create a single whole-slice Stage-3 chain from the weighted combination of the hemisphere-specific Stage-3 transformations. This ensures that the alignment remains precise in both hemispheres irrespective of variations in the interhemispheric gap or the antero-posterior shearing of the hemispheres (Figure 7G), which is particularly important for registering histology sections that were extracted from the midline. Finally, we load the histology data as a TImage and initialise it in MRI space by concatenating the optimised chains from Stages 1-2 and the whole-slice Stage-3 chain.

#### Registration

Stage 4 imports the so initialised histology image and the structural MRI volume. To reduce the complexity of the direct histology-to-MRI registration, a new Stage-4 chain is introduced that consists of the following transformations: a 3D displacement field (defined sparsely by a handful of control points, typically 16, scattered evenly across the histology domain), a 3D rotation (about the centre of the histology domain), and a 3D translation. The parameters of the new chain are set to provide a nearly equivalent mapping between histology and MRI space as the combined Stage 1-3 chain. By reducing the degrees of freedom of the non-linear transformation (from pixelwise in Stage 1 to 16 points in Stage 4), we reduce small-scale distortions of the histology image that have most likely arisen from anatomical disparities with the blockface photograph or the granularity of the histology stain. Finally, the parameters of the initialised Stage-4 chain are fine-tuned in a 4-step optimisation sequence that is similar to what was described for Stage 3. Here, the rigid search range is narrowed down, such that the histology section in MRI space is only allowed to travel ±2 mm perpendicularly to its initial orientation, accounting for the anatomical discrepancies that may be present as a result of sectioning the tissue block at greater depths.

#### Experiments

We ran Stage-4 optimisations on all 87 PLP-stained histology images that we had previously registered to MRI (TRUFI) data using Stages 1-3. The Stage-4 outcomes were visually compared with the Stage 1-3 outputs with special attention to the amount of distortions and the alignment of anatomical contours.

## 3. Results

### 3.1. Stage 1

The panels in Figure 8 show the registration error (MCD) for the linear and non-linear optimisation steps of the histology-to-block registration routine (Stage 1) for the 14 callosal and the 14 hippocampal sections. Actual registration results with different regularisation weights can be viewed in Supplementary material 4. Registration errors were also compared for different regularisation weights (a) in both anatomical regions. The linear registration steps constituted a gradual improvement in the alignment of the images. The non-linear substage significantly improved the accuracy of the registration, confirming the distorted state of the histology images. For the callosal sections, the difference between the MCD after the rotation search and the similarity transform was minimal, and the affine substage seemed to have a stronger influence on the images that were highly misaligned after the previous stages. An inspection of these images revealed that shears were interfering with the correct estimation of the rotation and the scale factor in the first substages, which were successfully compensated by the affine substage. The regularisation weight had an optimum at 0.3 <α < 0.4 (median MCD 0.22mm-0.23mm). At the highest regularisation value of a = 1.0 the alignment was still noticeably better compared to the result of the affine registration, especially for sections with larger initial misalignment (as evidenced by shrinkage of the interquartile ranges and the top bar). This is in keeping with the existence of bulk deformations (e.g., a slight bending of a gyrus) in the mounted tissue slice that cannot be compensated by global transformations but are still captured by the highly regularised non-linear substage. On the contrary, high regularisation cannot account for some finer distortions of tissue (e.g., anisotropic stretching). The pixelwise Jacobian determinants were positive across the images for all a > 0.2, indicating that no topological errors are induced by the registration.

**Figure 8.**
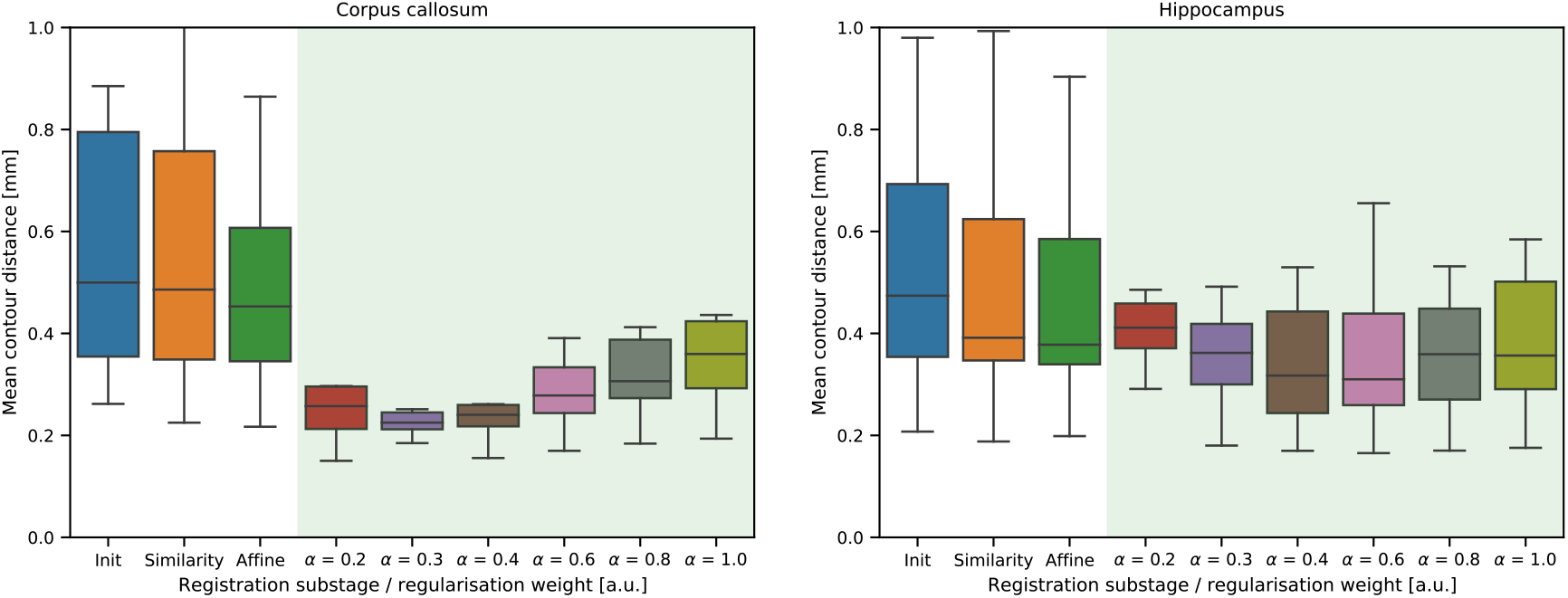
Registration error expressed as median contour distances (in mm) shown for 14 callosal and 14 hippocampal sections after the linear (*white background*) and non-linear (*green background*) steps of Stage 1. The non-linear registration error is reported for a range of different regularisation weights.

The registration of the same callosal sections with two different ANTs paradigms yielded consistently higher MCDs (SyN+Mattes: 0.4 mm, SyN+CC: 0.25 mm) than the results obtained by our Stage 1 routine (0.23 mm). The same trend was observed for hippocampal sections as well, although the registration errors were generally higher in this anatomical region (SyN+Mattes: 0.65 mm, SyN+CC: 0.6 mm, Stage 1: 0.4 mm). The distribution of MCDs (Figure 9) also reveals that the ANTs registration paradigms were generally less robust than the Stage-1 routine, with more frequent misregistrations in the affine stage. Supplementing the ANTs SyN+Mattes registration paradigm with TIRL-generated binary masks did not improve, rather aggravated affine initialisation errors in both anatomical regions. Representative registration results are shown in Figure 9. The runtimes for the two software were comparable: averaging around 1 minute for ANTs, and around 1 minute and 15 seconds for Stage 1.

**Figure 9.**
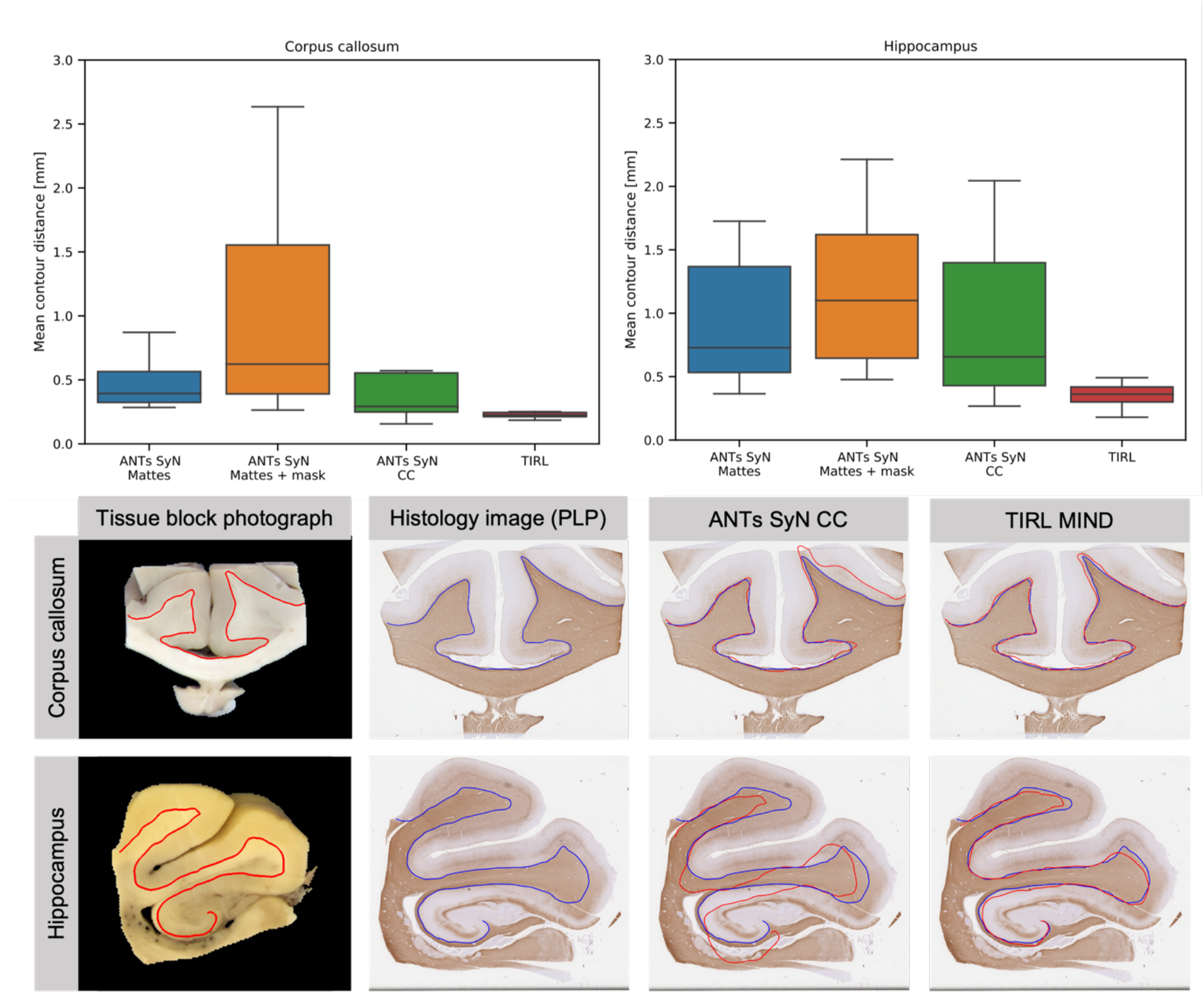
Comparison of histology-to-block registration by Stage 1 and various ANTs paradigms. *Top:* distribution of the registration error (MCDs in mm) corresponding to the four registration paradigms tested on 14 callosal and 14 hippocampal slides. *Bottom:* a visual comparison of registration results on representative callosal and hippocampal sections obtained with TIRL Stage 1 and ANTs SyN CC registration. The *red and blue contours* represent manual segmentations of the grey-white matter boundary in the tissue block photo and the PLP-stained histology images, respectively. These and similar contours were used to compute the MCDs.

### 3.2. Stage 2

Upon careful observation, 81 out of 87 block-to-slice registrations were highly accurate: grey-white boundaries were well-aligned and characteristic small features of the images, such as penetrating vascular structures, were generally seen within a 4-pixel range (<0.2 mm) from each other, which in MRI terms translates to sub-voxel precision even for our high-resolution TRUFI data (0.25 mm/voxel). Figure 10 shows representative registration results from various anatomical regions. MCD measurements on 12 randomly chosen examples confirmed the <0.2mm accuracy. In the remaining 6 out of 87 cases, the registration could not succeed due to some form of human error, such as incorrect labelling of the slice photograph or the tissue block, misidentification of the corresponding slice, or block surface. After fixing these, Stage 2 yielded equally accurate results for these images as well.

**Figure 10.**
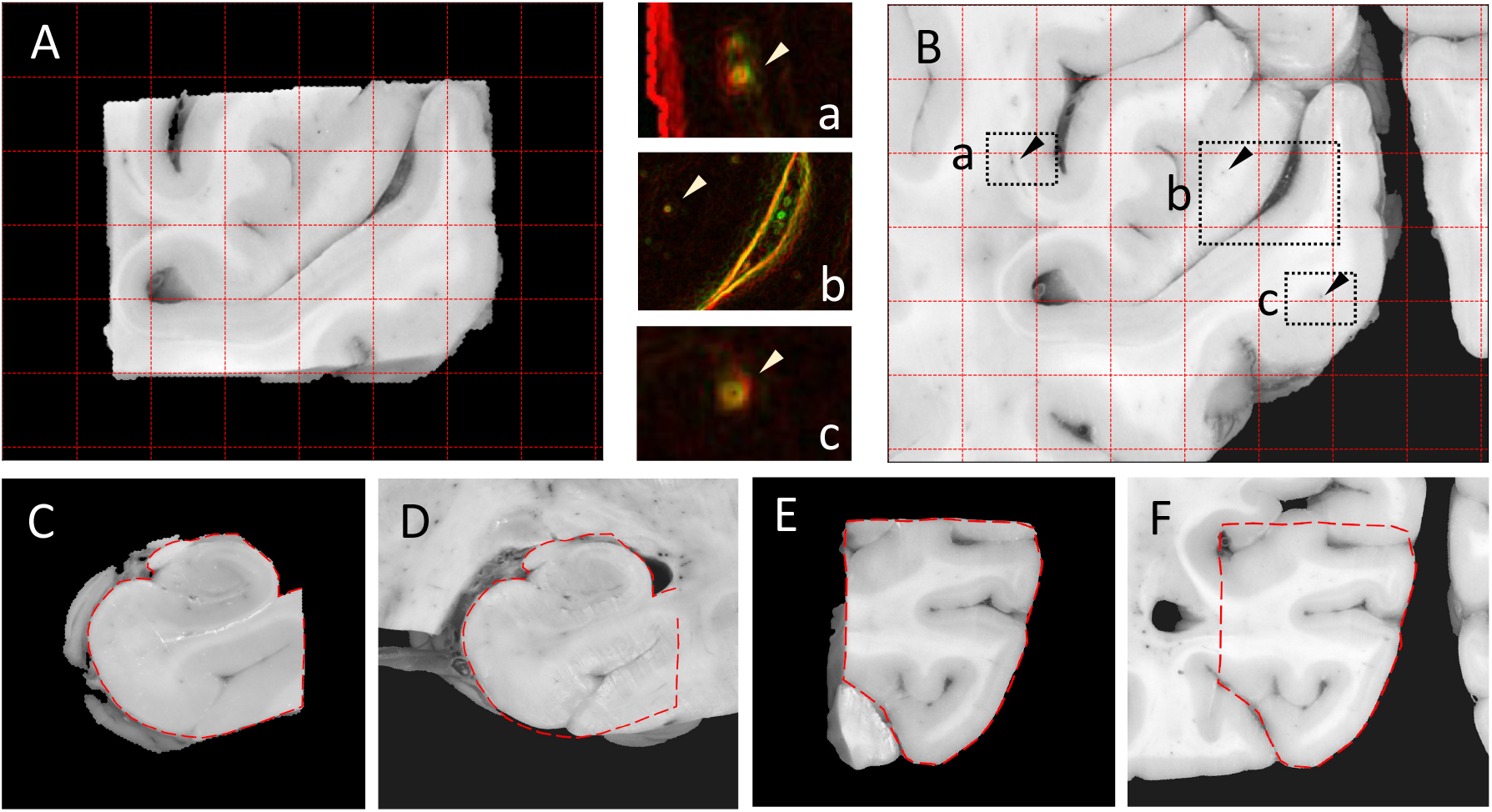
Accuracy of Stage-2 registration of tissue blocks in various anatomical regions. **(A)** Tissue block photograph showing the left visual cortex. Grid spacing: 5 mm. **(B)** Left visual cortex region of the corresponding brain slice photograph shown after alignment with (A). **(a,b,c)**: colour-coded edge-enhanced overlay of (A, red) and (B, green) within the marked regions, demonstrating the alignment of perforating vessels. The yellow colour emerges from red-green overlap, indicating accurate alignment between anatomical contours. **(C-D)** Registered right hippocampus block. **(E-F)** Registered left parahippocampal gyrus.

In our robustness experiment, the automatic block initialisation routine could successfully identify as many as 8 different sampling sites on the same brain slice, and all corresponding tissue blocks could be assigned to the correct sampling site. Comparing the initial and final positions of all 87 blocks at their respective sampling sites, we further found that the error of the automatic block initialisation routine was consistently low, with a median value of 0.49 mm, and a 95^th^ percentile of 2.54 mm.

The resilience of the registration algorithm against initialisation errors is vital for robust performance at Stage 2. To test this resilience, we simulated 100 different initialisations for each of the previously mentioned 8 blocks and recorded whether they led to a successful (MCD < 0.2 mm) or an unsuccessful registration of the blocks. Figure 11 shows that the overwhelming majority of the simulated registrations were successful for all blocks, and most unsuccessful registrations occurred when the blocks were initialised far away from the centre of the sampling site. Importantly, no failures were observed within the median initialisation error mark for any of the blocks. Two failures were observed for blocks *b* and *c*, and one for block *g* within the 95^th^ percentile radius. For blocks with less salient anatomical features (b, c, d) there was a marked decrease in the success rate (Figure 11, plots) beyond the 95^th^ percentile error mark, which only occurred after 7.5mm for the blocks that presented with clear contrast.

**Figure 11.**
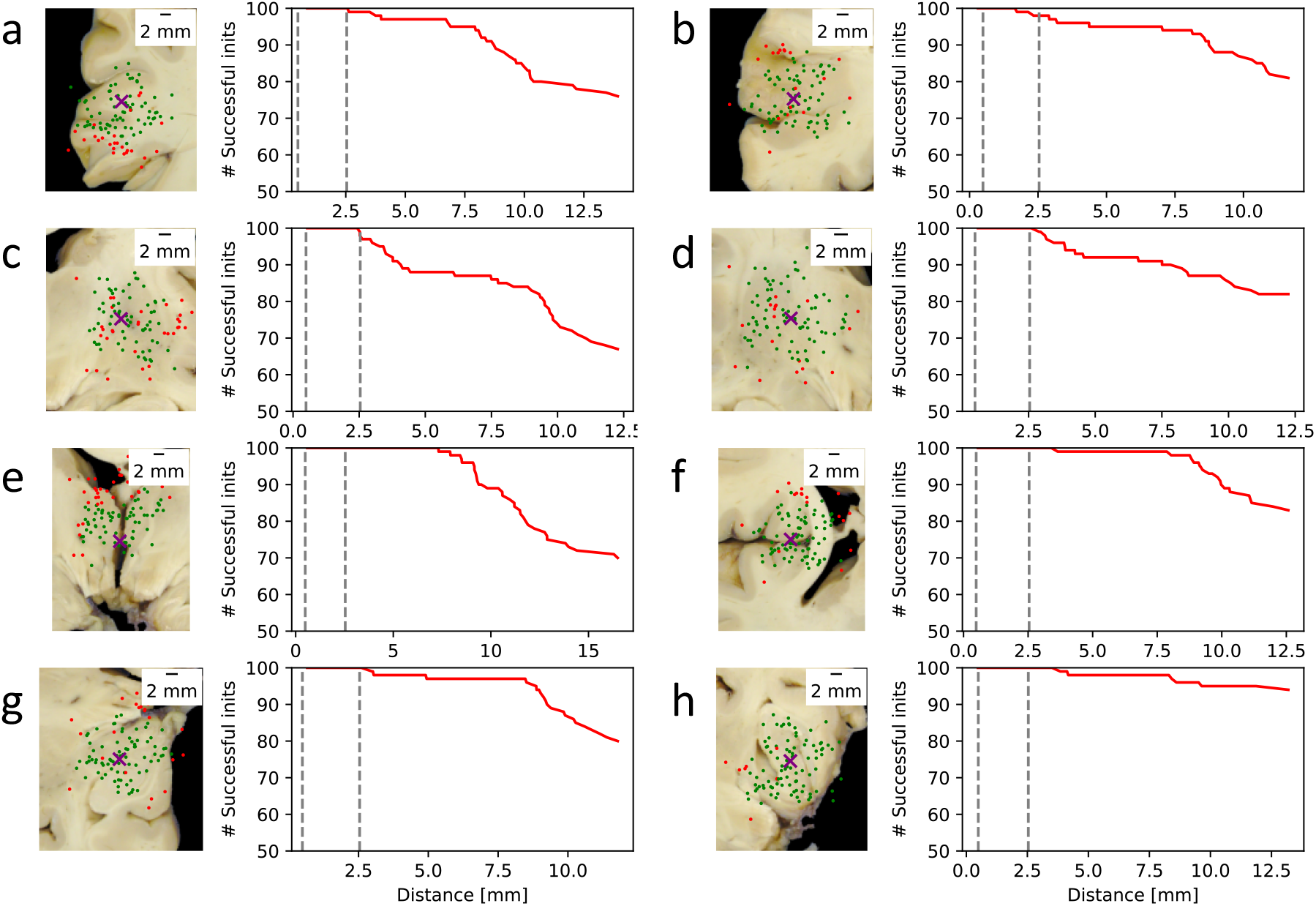
Resilience of the Stage-2 registration algorithm against simulated block initialisation errors. The *purple crosses* mark the true centre of the blocks on the brain slice photographs. The *green dots* represent random initialisations associated with a successful (MCD < 0.2 mm) registration result, whereas *red dots* correspond to unsuccessful registrations. The *red graphs* represent 100 – #failures within a given radius from the true centre. The *dashed grey lines* indicate the median and the 95^th^ percentile values of the true block initialisation error measured on the whole dataset.

The results of the robustness experiment indicate that with the default configurations, Stage-2 block registrations are highly accurate (<0.2mm). Failures can be more prevalent for blocks with less salient anatomical features, but only if the insertion site is offset by more than 2.5mm. If a misregistration occurs due to erroneous initialisation, manual intervention is needed to provide a more appropriate initial position for a block. Alternatively, the grid search of the rotation and translation parameters may be expanded to attempt to increase the robustness of the pipeline, though this will be at the expense of additional computational cost.

### 3.3. Stage 3

#### 3.3.1. Registration results with actual brain slice photographs

Figure 12 shows an example brain slice registration. The cutting plane was oblique, as evidenced by the asymmetric appearance of the lateral ventricles, the hemispheres were completely detached, and the cerebellum was not represented in the photograph (Figure 12A). Similar phenomena were found to be common across the dataset. Stage 3 was run with default configurations, and the accuracy of the registration was assessed qualitatively by overlaying grey-white matter contours after each registration step. The rigid and affine registration steps estimated the obliqueness of the cut surface accurately enough to reproduce the gross shape of the hemispherical cross sections with the asymmetric appearance of the lateral ventricles (Figure 12B). However, a closer inspection of the reconstructed MRI slice with the contours reveals several regions where the affine registration was less accurate (yellow arrowheads in Figure 12B). In most of these regions, the contours are not only misaligned but anatomically different – a hallmark that the slicing plane could not be fully estimated by the affine transformation, most likely because it is curved. Accordingly, the misalignments in these regions persisted after optimising in-plane deformations (Figure 12C). On the contrary, the free-form deformation step could achieve an almost perfect alignment of the grey-white contours (Figure 12D). The presented final registration accuracy is representative of all 209 slices that we processed with Stage 3. Generally, the slices were registered with through-plane deformations exceeding the MRI voxel size, and in this specific case, they were as large as 6.5mm. The Jacobian exhibited a conservative ±10% dilation/shrinkage of the slice throughout the entire 2D/3D transformation (Figure 12E), indicating the non-linear deformations were predominantly related to slice curvature. The spatial distribution of in-plane and through-plane deformations appears to be consistent with a combination of two factors: the bulk deformations of the brain while it is loaded into a plastic mould for scanning, as well as the compression and shearing of the hemispheres while the brain is cut (Figure 12F).

**Figure 12.**
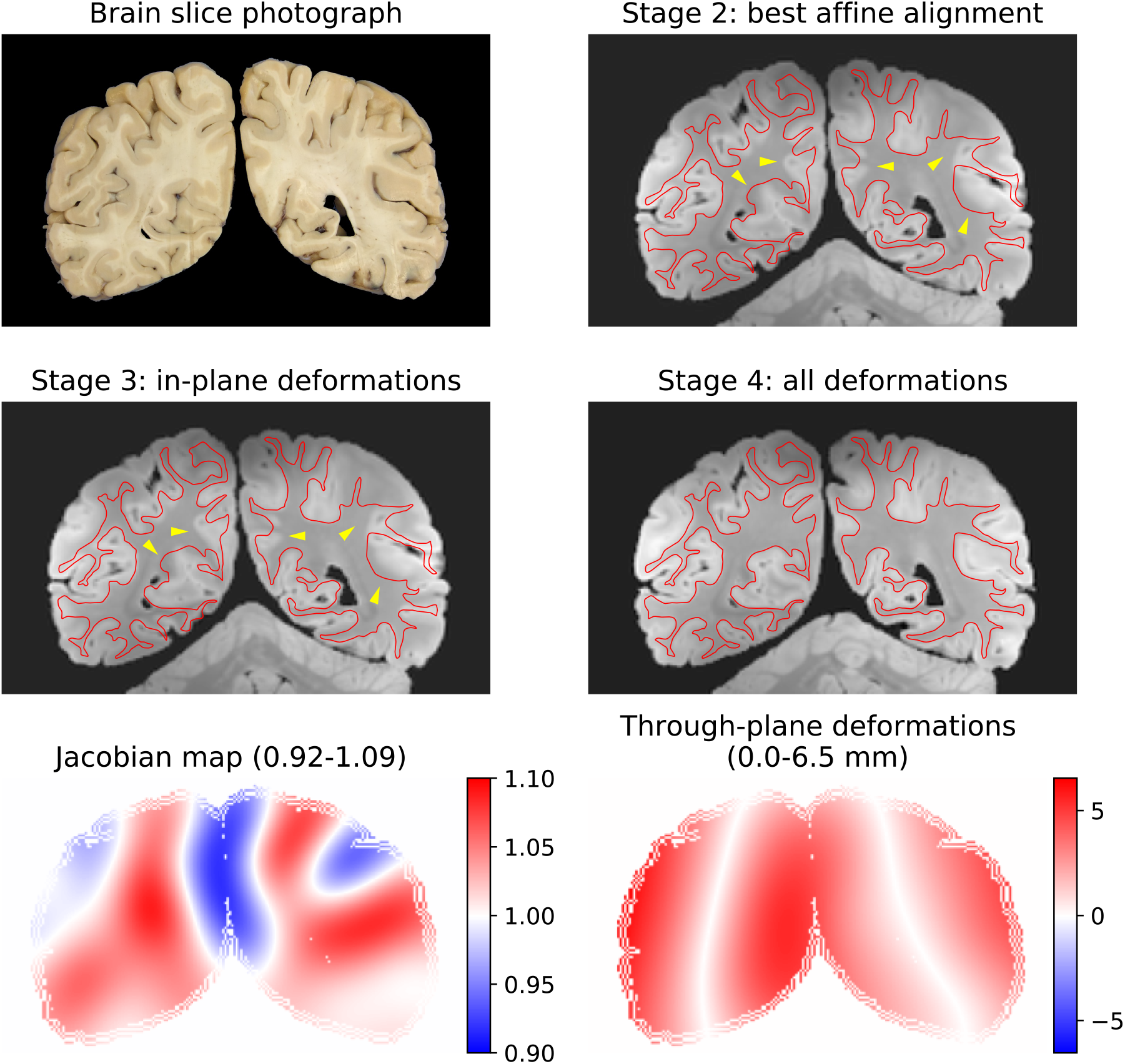
Stage-3 (slice-to-volume) registration result of an actual coronal brain slice photograph. The grey-white matter boundary (*red contour*) was segmented by hand on the brain slice photograph **(A)** and overlaid on the resampled MRI data after each optimisation step **(B-D)** to assess the accuracy of the registration. Notable misalignments are indicated by the *yellow arrowheads*. Through-plane deformations **(D)** are essential for an accurate registration of this slice. **(E)** The conservative range of Jacobians suggest moderate in-plane deformations, while the 3D deformations of the slicing plane **(F)** are remarkable (the scale shows displacements in mm).

#### 3.3.2. Registration results with simulated brain slices

Using simulated slices, which were generated by resampling the structural MRI data of a single subject onto a series of analytically defined surfaces (Figure 13), we could further quantify the registration accuracy and robustness. Starting from a perturbed position and orientation, all slices registered well with no exceptions, and the MRE showed a steady decline across consecutive optimisation steps (Table 1).

**Figure 13.**
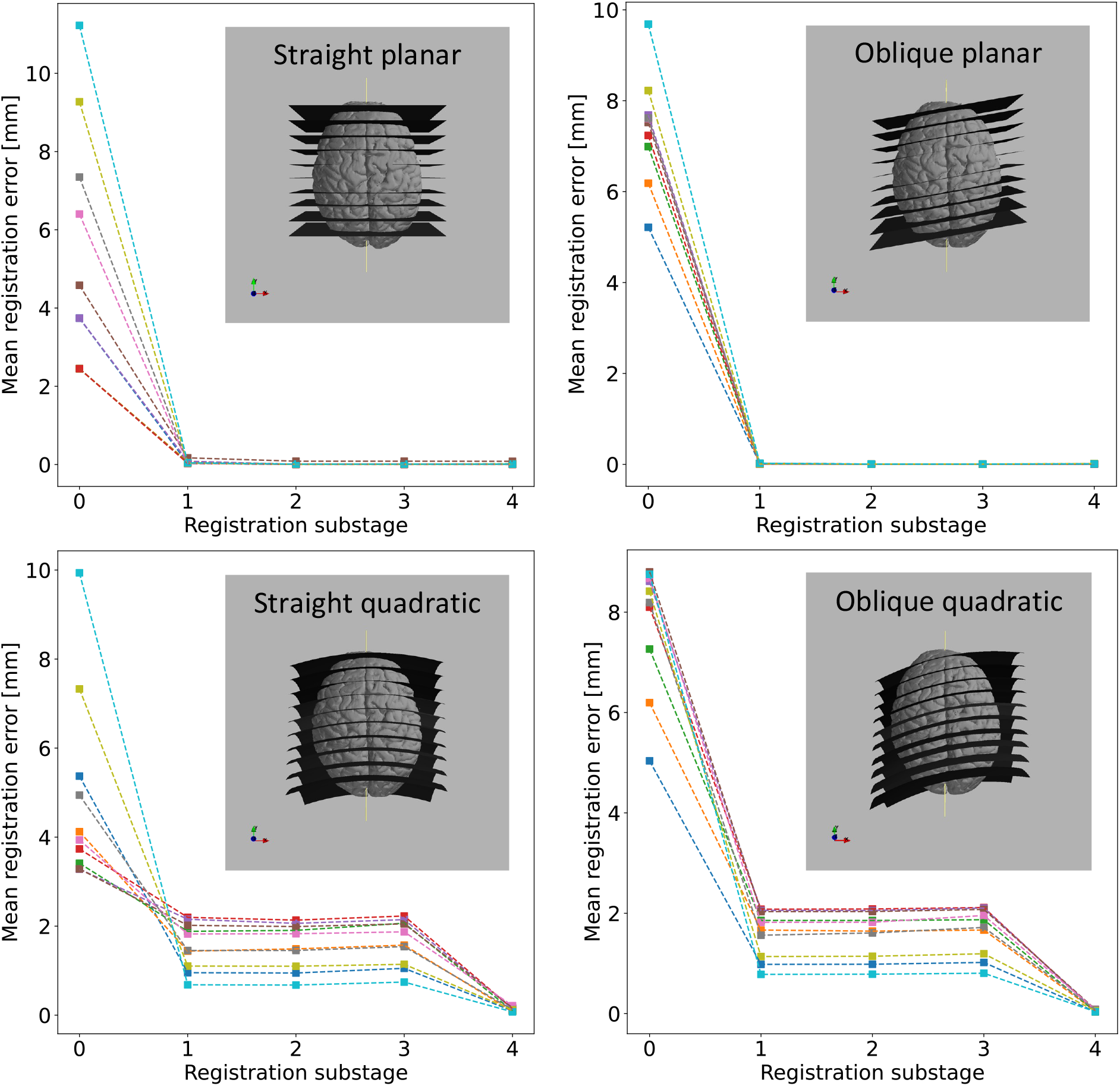
Quantifying Stage-3 (slice-to-volume) registration error using four different sets of simulated slices. Each series (straight planar, oblique planar, straight quadratic, oblique quadratic) consists of 10 simulated slices in the postero-anterior direction. The median registration error (MRE) is plotted for each slice after each optimisation step (0: perturbed initial state, 1: rigid, 2: affine, 3: in-plane deformations, 4: 3D deformations). The gradual convergence of the MRE towards zero in all cases demonstrates the robustness of Stage 3 as well as the added value of each optimisation step.

**Table 1.** Median slice registration error (MRE, in mm) after each optimisation step of Stage 3. Step 0 refers to the initial state, and the numbers represent the extent of slice perturbations. The optimisation steps are 1: rigid, 2: affine, 3: in-plane deformations, 4: 3D deformations. The output of step 4 (highlighted) was accepted as the final output from Stage 3 in all cases.

| Step | Slice series |  |  |  |
| --- | --- | --- | --- | --- |
|  | Straight | Oblique | Straight quadratic | Oblique quadratic |
| 0 | 5.366 | 7.397 | 4.935 | 7.805 |
| 1 | 0.058 | 0.012 | 1.570 | 1.611 |
| 2 | 0.013 | 0.004 | 1.558 | 1.604 |
| 3 | 0.013 | 0.004 | 1.641 | 1.667 |
| 4 | 0.015 | 0.008 | 0.125 | 0.126 |

##### Planar slices

Following the initial perturbation of the 10 straight slices (Table 1, column 1), the MREs were uniformly distributed between 2.5 mm and 12 mm with an average MRE of 5.36 mm. Despite the large variation, all 10 slices could be registered equally well by the rigid substage (MRE = 0.058 mm), which was further improved to MRE = 0.014 mm by the affine optimisation. The MRE stayed roughly constant through the non-linear optimisation steps. As for the oblique slices (Table 1, column 2), despite the larger initial perturbations (average MRE = 7.397 mm), the rigid optimisation step successfully registered all 10 slices (MRE = 0.013 mm), and the affine optimisation made a further improvement (MRE = 0.004 mm), which stayed roughly constant during the non-linear steps. These results underpin that the linear optimisation steps converge to the desired optimum, regardless of whether the initialisation is close or further away, and that Stage 3 does not introduce unnecessary slice deformations.

##### Quadratic slices

The straight (Table 1, column 3) and oblique series (Table 1, column 4) showed a common trend of the MRE, but this was qualitatively different from that of the planar sections (Figure 13). The random perturbations were somewhat larger for the oblique series (MRE: 4.94 mm vs. 7.81 mm), but this difference completely vanished after the rigid substage (MRE: 1.57 mm vs. 1.61 mm), indicating that the rotation components were accurately estimated for the oblique slices despite their coexisting curvature. The MREs after the rigid alignment were comparable in size to the deflections of the planes (<3 mm), and neither the affine transformation (MRE: 1.56 mm vs. 1.60 mm), nor the in-plane deformations (MRE: 1.64 mm vs. 1.66 mm) could make any improvement. In fact, the in-plane optimisation slightly increased the registration error by falsely attributing slice curvature related misalignments to in-plane deformations. These were reverted and successfully converted to orthogonal displacements in the 4^th^ optimisation step, as indicated by the MREs of 0.125 mm and 0.126 mm for the straight and oblique series, respectively. This result demonstrates that Stage 3 can converge on physically realistic curvatures of the brain slices from an initial planar estimate, and the curvatures can be estimated with sub-voxel (<0.25 mm) precision.

### 3.4. Stage 4

We observed significant improvements in the alignment of histology and MRI features after the Stage-4 optimisation of Stage 1-3 results, for sections where the primary source of error was the anatomical discrepancy between the histology and the tissue block photograph (Figure 14). Stage 4 also dramatically improved the registration accuracy for histology sections that were sampled from across the interhemispheric fissure (e.g., anterior cingulate cortices, corpus callosum). Regions with limited anatomical features, however, struggled to drive the non-linear steps of Stage 4, and often led to exaggerated deformations. This led us to conclude that the linear optimisation steps of Stage 4 provided the best overall match between PLP and TRUFI data in our dataset, but further steps can easily be specified in the Stage 4 configurations for other datasets.

**Figure 14.**
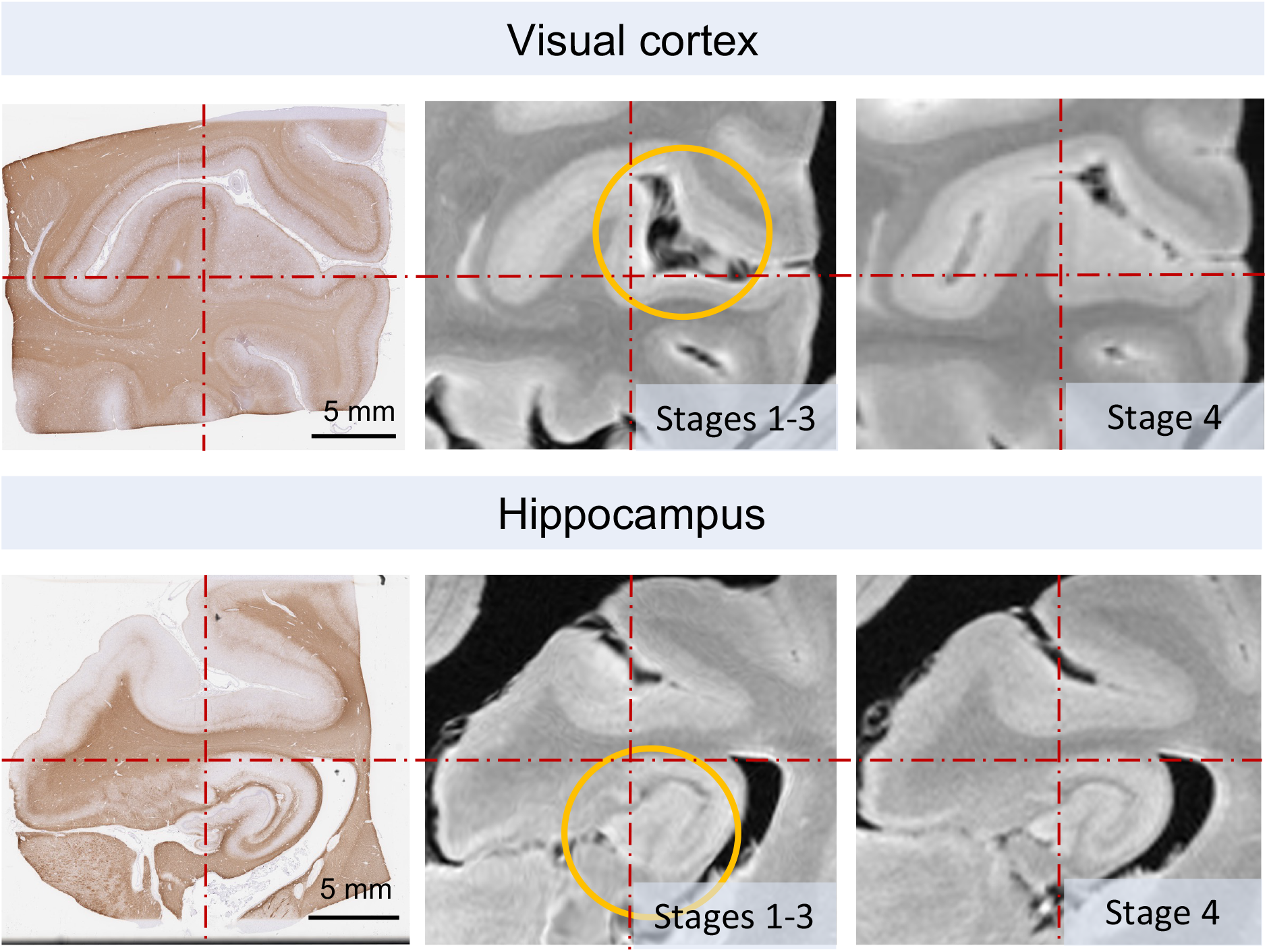
Stage 4 can improve the accuracy of histology-to-MRI registration. *Top row:* visual cortex, *Bottom row:* hippocampus. In both cases, the histology sections were sampled from deeper inside the tissue block, hence they exhibit a slightly different anatomical pattern than the corresponding tissue block photographs that were used in Stage 1. The *red centre lines* are provided to guide the eye. The main areas of improvement after Stage 4 are highlighted by the *orange circles*.Also note that tissue contours appear less distorted in the Stage 4 results, because Stage 4 deformations are defined with fewer degrees of freedom to mitigate any previously overestimated deformations of the tissue.

We resampled multi-modal MRI data onto the registered histology domain by concatenating and applying the Stage-4 optimised chains with the respective inter-modality linear transformation matrices obtained from FLIRT. The MRI modalities included both scalar and vector quantities, the latter of which were adequately rotated in 3D by the combined transformation chain. Using an adapted version of the Stage 1 script (available via Git) we could also register histology sections of various stains onto the domain of the respective PLP-stained section. Since this was originally used for the registration with MRI, we could achieve a pixel-to-voxel mapping between any pair of histology and MRI modality by concatenating the histology-to-histology and histology-to-MRI chains (Figure 15).

**Figure 15.**
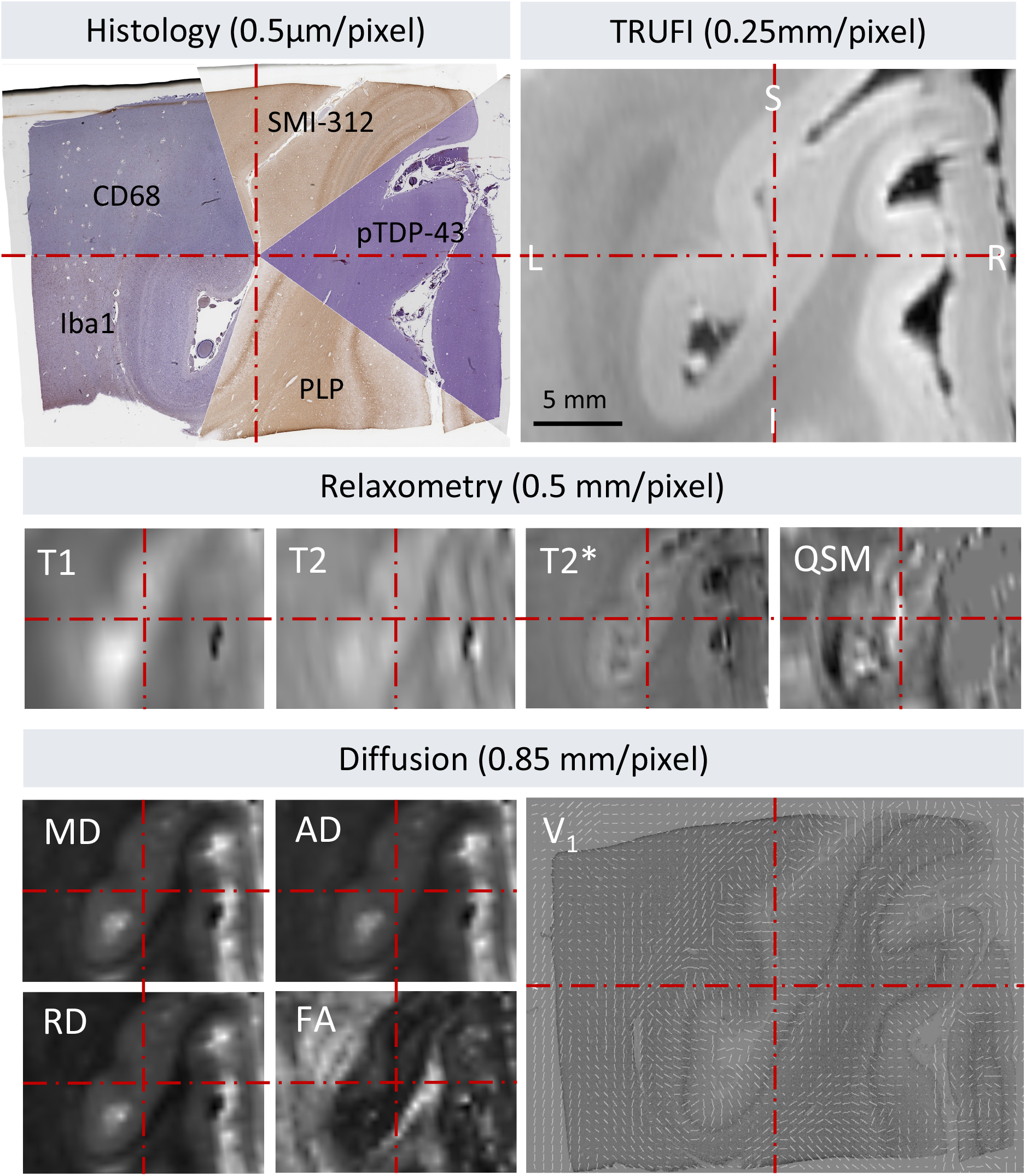
Example of a registered MRI-histology stack of the left visual cortex, consisting of five histology stains (PLP, Iba1, CD68, SMI-312, pTDP-43), various relaxometry (T1, T2, T2*, QSM), and diffusion MRI modalities (MD, AD, RD, FA, V_1_). All images are pixelwise aligned (the *red centre lines* are provided to guide the eye). **V_1_**: PLP-stained histological section of the left visual cortex overlaid with a map of principal fibre orientations derived from post-mortem diffusion MRI data via diffusion tensor fitting. The fibre orientation vectors are automatically rotated by TIRL in accordance with the transformations of the histology slide.

## 4. Discussion

In this paper, we presented a novel image registration framework, TIRL, and used it to create an automated histology-to-MRI registration pipeline that was specifically designed to work with sparsely sampled histology data, unlike most existing methods that require serial histological sectioning. Nevertheless, there is nothing to prevent one from registering adjacent sections of densely sampled histology data to MRI using TIRL, as we recently demonstrated in a macaque brain dataset with preliminary results [59, 60].

Our results with deformable 2D-to-3D registration demonstrate the value of compensating for 3D slice deformations to achieve submillimetre accurate alignment between histology and MRI. Our method does not rely on specialised cutting or stain automation hardware for tissue processing and reduces the imperfections of alignment that arise from freehand brain cutting, making it suitable for integration into routine neuropathological practice. The four stages of the pipeline support full automation of the registration, provided that the relevant dissection photographs are available. With suitable manual initialisation however, Stage 4 could also be used as a stand-alone semi-automated tool in the absence of photos to directly register histology images to volumetric MRI data, although in this work we have not tested the performance of this specific usage.

TIRL was designed to be a general image registration tool that can support a wide range of applications, which could include a range of species, organ systems, and pathologies. The implementation of the MIND cost function [55] in TIRL ensures that our pipeline is compatible with a diverse range of image modalities, and the modular implementation of the library allows further intuitive and straightforward customisation of its components.

In our efforts to make TIRL as general as possible, we had to make occasional trade-offs with computational efficiency. Nevertheless, we made significant efforts to include computational optimisations where possible, such as parallel processing, chunked interpolation, function caching, optimising sub-chains of linear transformations by affine replacement, and avoiding interpolation of displacement fields where the field is defined over the same domain as the image. Our experiments were carried out on a Dell T7500 workstation computer with two hexa-core Intel X5670 CPUs (2.93 GHz) and 64 GB of RAM. The typical runtimes were ^~^2 minutes for stage 1, ^~^30 minutes for stage 2 (with 6 insertion sites), 1–2 hours for stage 3 (using 50 control points), and ^~^15 minutes for stage 4. For relatively undistorted slices, it is possible to reduce the runtime of stage 3 by using fewer control points (≤16) to optimise deformations. Stages 1-3 can be run in parallel, while Stage 4 requires the outputs of the earlier stages.

Based on our personal experience, future users should observe the following data acquisition principles to achieve high-quality registration results with our pipeline:

1. **Histology** sections should be sampled as close as possible (<1 mm) to the photographed surface of the tissue blocks. Care should be taken to avoid tears and folds of the tissue as well as staining artefacts during the histological processing. At least one stain with sufficient anatomical contrast (in brain, grey-white matter) must be available for registration with MRI. This specific stain can then guide the alignment of other stains without this contrast.
2. **Dissection:** Standardising the position and orientation of the brain cuts makes it easier to initialise slice-to-volume registrations (Stage 3).
3. **Photographs:** Photographs should be taken at high resolution, under diffuse lighting conditions, on a clean, matte surface. The background should have a distinct colour from the brain tissue to allow segmentation. Brain slices should be photographed on both sides, avoiding glare from any lighting. The approximate mm/pixel resolution of the photographs should be recorded. The slices should be identified with labels within the photographs to avoid mix-up.
4. **MRI:** MRI should be acquired at high resolution (0.25–1 mm/voxel) with strong contrast of relevant anatomy. Specimens should ideally be scanned in a container that is tailored to the shape of the specimens to avoid excessive deformations (small deformations can be corrected by the pipeline). The container should be filled with a susceptibility-matched, signal-free fluid (e.g., perfluorocarbon such as Fluorinert) [61], and air bubbles should be avoided [54].

We have committed to sharing our multi-modal MRI-histology data via the Oxford Digital Brain Bank (https://open.win.ox.ac.uk/DigitalBrainBank/#/datasets/pathologist). Both TIRL and the MRI-histology registration pipeline are distributed in the form of Git repositories, and as part of FSL (v6.0.4 and above). We hope that this will facilitate MRI-histology research and encourage the development of further analysis tools built on top of TIRL, paving the way toward more histologically validated imaging studies in the future.

## 5. Conclusion

A novel image registration framework, TIRL was presented through its application to create an automated pipeline for registering sparsely sampled histology sections to volumetric post-mortem MRI data. The pipeline accounts for 3D deformations of thin tissue sections, does not require manual intervention in most cases, and achieves submillimetre registration accuracy through photographic intermediaries, which can be readily acquired as part of routine neuropathological practice. The customisability of the pipeline and the underlying software framework present a great appeal for future histology-MRI investigations.

## Author contributions

- I.N. Huszar: Designed, implemented, tested TIRL and all scripts of the registration pipeline, created figures, wrote and edited manuscript.
- M. Pallebage-Gamarallage: Designed the histopathological protocol of the MND study, dissected brains, took photographs and created stained histological specimens, edited manuscript.
- S. Bangerter-Christensen: Prepared various stained histological specimens, edited manuscript.
- H. Brooks: Prepared LFB-stained histological specimens, edited manuscript.
- S. Fitzgibbon: Tested and provided feedback about the implementation of Stage 1 and the TIRL framework, edited manuscript.
- S. Foxley: Designed the post-mortem MRI protocol of the MND study and acquired MRI data, edited manuscript.
- M. Hiemstra: Prepared stained histological specimens of the hippocampus, provided feedback about Stage 1, edited manuscript.
- A.F.D. Howard: Tested and provided feedback about the implementation of Stage 3 and the TIRL framework, edited manuscript.
- S. Jbabdi: Tested and provided feedback about the implementation of Stage 1 and the TIRL framework, edited manuscript.
- D.Z.L. Kor: Tested the TIRL framework, provided feedback about the Stage 4, edited manuscript.
- A. Leonte: Prepared stained histological specimens of the anterior cingulate cortex, edited manuscript.
- J. Mollink: Prepared stained histological specimens of the hippocampus, provided feedback about Stage 1, edited manuscript.
- A. Smart: Prepared various stained histological specimens, edited manuscript.
- B. C. Tendler: Created post-processing pipeline for post-mortem MRI data, provided feedback about Stage 4, edited manuscript.
- M. R. Turner: Designed MND study, provided neurological expertise, edited manuscript.
- O. Ansorge: Designed MND study, provided neuropathological expertise, and material from the Oxford Brain Bank, obtained funding, edited manuscript.
- K. L. Miller: Designed MND study, provided MRI physics expertise, obtained funding, edited manuscript.
- M. Jenkinson: Provided image analysis expertise, designed TIRL, the registration pipeline and the experiments, edited manuscript.

## Acknowledgements

The authors express their gratitude to the donors and benefactors of the Oxford Brain Bank, that kindly provided all human tissues for this study. Core funding for the Oxford Brain Bank was provided by the Medical Research Council (MRC), the NIHR Oxford Biomedical Research Centre and the Brains for Dementia Research programme, jointly funded by Alzheimer’s Research UK and Alzheimer’s Society, in association with the MRC. INH and AFDH were supported by the Engineering and Physical Sciences Research Council (EPSRC, EP/L016052/1) and the Medical Research Council (MRC, MR/L009013/1) and the Wellcome Trust (WT202788/Z/16/A). INH received additional funding from the Clarendon Fund in partnership with the Chadwyck-Healey Charitable Trust at Kellogg College (Oxford). MJ and OA were supported by the National Institute for Health Research (NIHR) Oxford Biomedical Research Centre (BRC). MPG, SF and the dataset used in this study were funded by an MRC Project Grant (MR/K02213X/1). KLM, BCT and JM were funded by a Wellcome Trust Senior Research Fellowship (202788/Z/16/Z). MRT is supported by a grant from the Motor Neurone Disease Association. The Wellcome Trust provided core funding for the Wellcome Centre for Integrative Neuroimaging (203139/Z/16/Z).

## Supplementary Material 1

### Details of the ANTs registrations that were used in the Stage-1 accuracy comparison testing

Comparisons were made with both the Mattes mutual information and the cross-correlation metrics, that were used in a previous study [25] to register histology sections. The registrations were carried out on the same 14 callosal and 14 hippocampal sections that were previously registered to their corresponding tissue blocks with the Stage 1 routine. For a fair comparison, all ANTs registrations were initialised using `antsAffineinitializer` with 30° increments over the whole circle. Various combinations of parameters were screened for each metric, starting from those that were recommended in the ANTs documentation. Empirically the following multi-resolution configurations were found to yield the best results with ANTs:

- ANTs SyN Mattes: antsRegistration -dimensionality 2 -float 0 -output $outdir/ants.syn/moving_to_fixed –interpolation Linear –winsorize-imageintensities [0.005,0.995] –use-histogram-matching 1 –r $outdir/ants.syn/init.mat -m Mattes[$outdir/fixed.png, $outdir/moving.png,1, 20, Random, 0.2] -t affine[2.0] –c [1500 × 1500 × 1500 × 300 × 100 × 0, 1.e-7, 5] –s 5×4×3×2×1×0 –f 7×6×5×4×2×1 –m Mattes[$outdir/fixed.png, $outdir/moving.png, 1, 32] –t syn[0.25,3.0,1] –c [200 × 200 × 200 × 200 × 150 × 50, 0, 5] –s 5×4×3×2×1×0 ×f 7×6×5×4×2×1
- ANTs SyN CC: antsRegistration –dimensionality 2 -float 0 –output $outdir/ants.syn.cc/moving_to_fixed –interpolation Linear –winsorize-image-intensities [0.005,0.995] –use-histogram-matching 1 –r $outdir/ants.syn.cc/init.mat –m Mattes[$outdir/fixed.png, $outdir/moving.png, 1, 20, Random, 0.2] –t

As the TIRL Stage 1 routine uses binary masks for the registration, the masks were exported from the TIRL pipeline, and the ANTs registrations were repeated with the masks. The previously generated contours of the tissue block photographs were transformed to histology space with the `antsApplyTransformsToPoints` tool, and the MCDs were calculated to measure the accuracy of the registrations in each case:

~~~
antsApplyTransformsToPoints –dimensionality 2 –precision 1 –input blockpts – –outputd/transformed_block_contour.csv –t [$d/moving_to_fixed0GenericAffine.mat,1] –t $d/moving_to_fixed1InverseWarp.nii.gz
~~~

## Supplementary Material 2

### Example Stage-3 registration of a severely damaged coronal brain slice using a manually defined binary mask for cost-function weighting

**Figure B.1.**
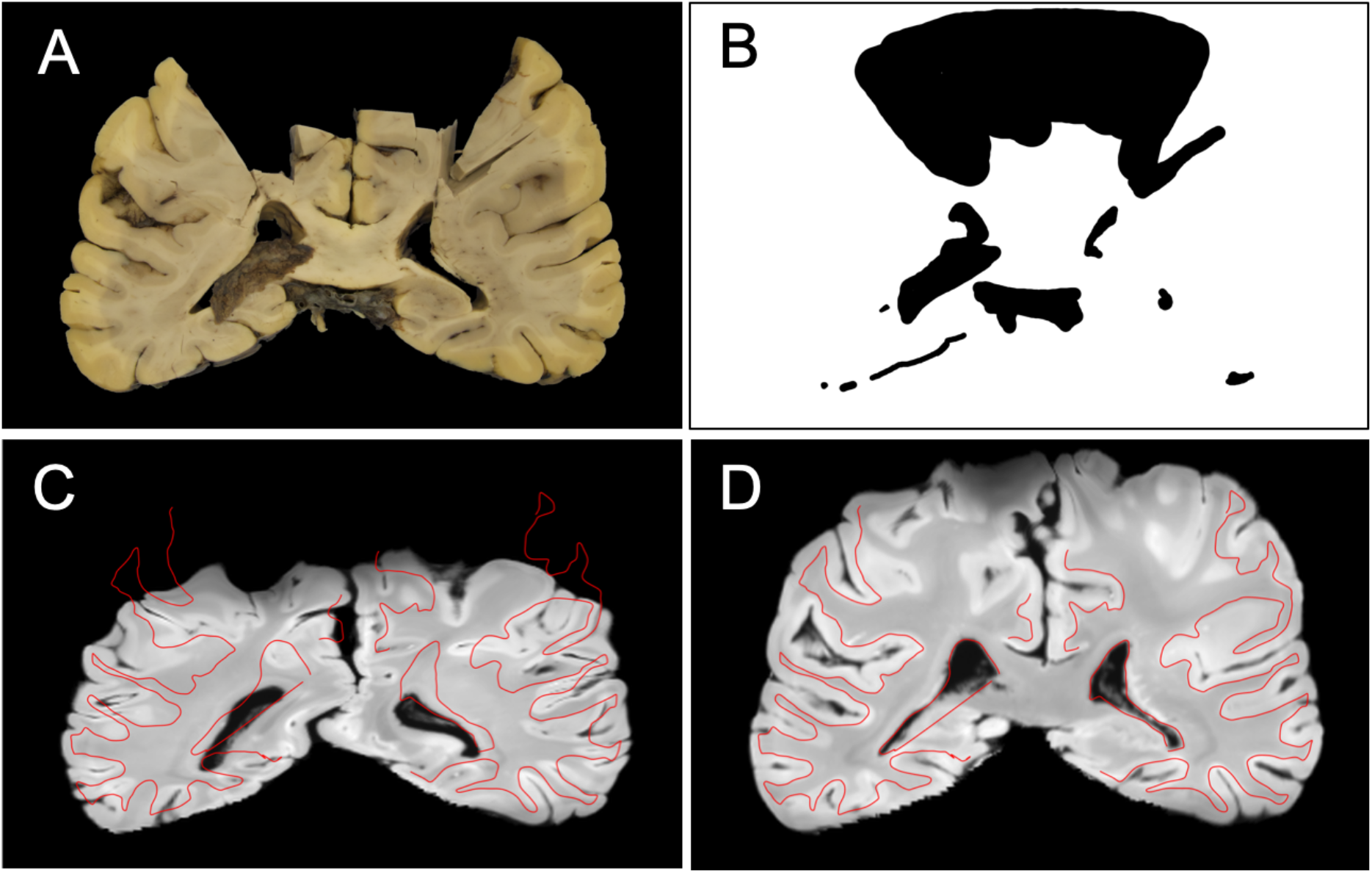
Result of slice-to-volume registration of a severely damaged coronal brain slice. **(A)** Coronal brain slice photograph with bilateral hiatus in the sensorimotor regions. **(B)** A hand-drawn binary mask for cost-function weighting. **(C)** Registration result without using the target mask. The *red curve* is an overlay of the manually segmented grey-white matter contour of the brain slice photograph. **(D)** Registration result with the hand-drawn target mask. The accuracy of the corrected registration is qualitatively similar to that on non-damaged slices.

## Supplementary Material 3

### Stage-3 simulation experiment with MIND and NMI cost functions

Figure C.1 shows that for oblique quadratic simulated slices with no Gaussian noise, MIND already outperforms NMI with respect to the final registration error (substage 4), which becomes even more obvious when Gaussian noise is added, mimicking the conditions of registering a slice of a different modality (e.g., a photograph). Based on the result of this experiment, NMI was dropped from the Stage-3 routine despite faster computations versus MIND.

**Figure C.1.**
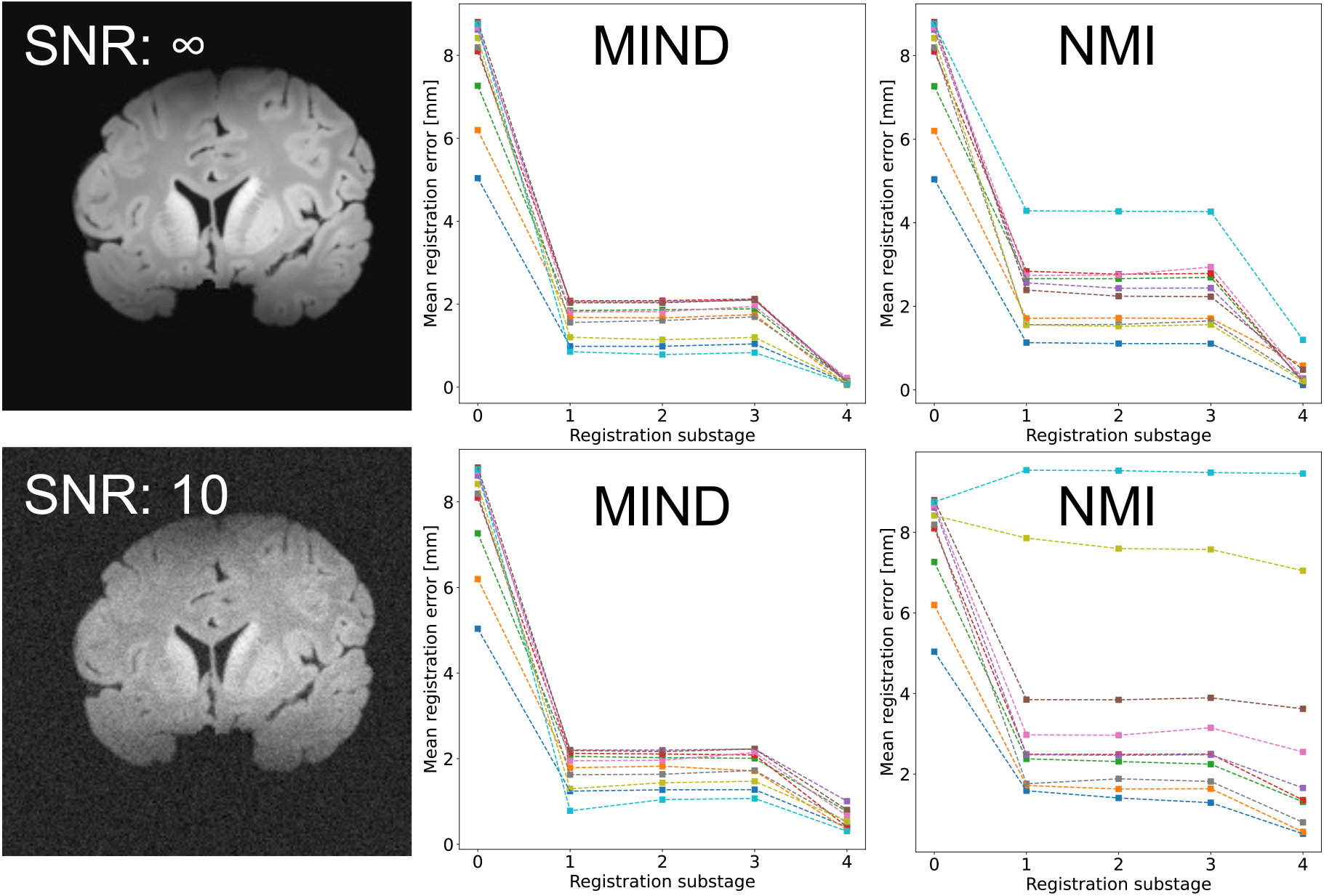
Comparison of the MIND and the normalised mutual information (NMI) image dissimilarity metric for Stage-3 registration of simulated slices (oblique quadratic series, using 16 control points in Steps 3 and 4). The registration substages are as described in the Stage-3 algorithm: 0) perturbed initial state, 1) rigid, 2) affine, 3) in-plane deformation, 4) 3D deformation.

## Supplementary Material 4

### Effect of regularisation weight on Stage-1 registration results: a visual comparison

**Figure D.1.**
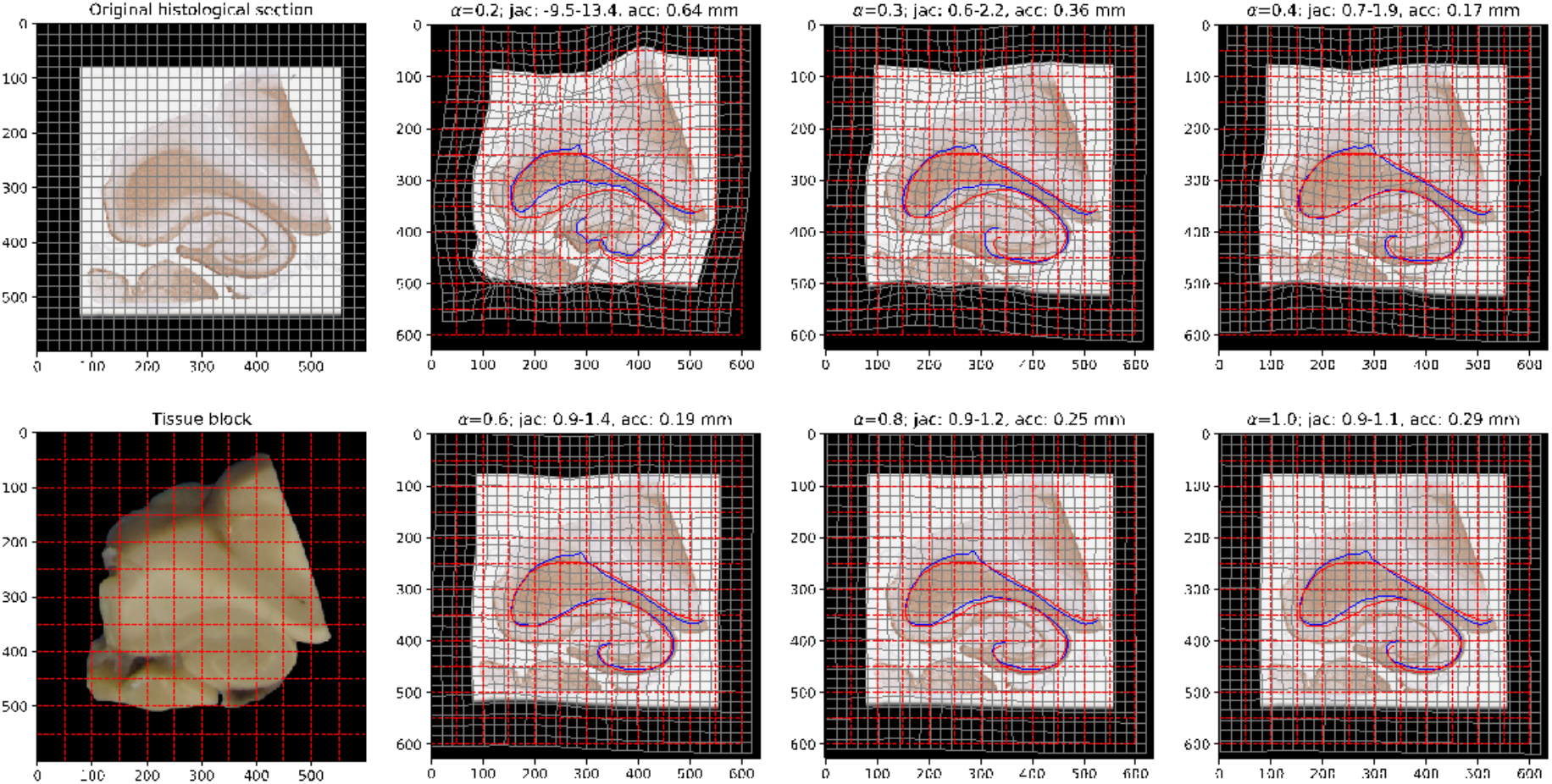
Stage-1 registration results on a hippocampal section using varying weights (0.2 < α < 1.0) for diffusion regularisation.

Images show deformed versions of the histology image on the domain of the tissue block photograph after the registration. The *blue curve* represents the transformed grey-white matter contour of the histology image, and the *red curve* is the grey-white matter boundary as observed in the tissue block photo. Their median distances are reported in millimetres above the images (*acc*). The Jacobian range (*jac*) is calculated from the total deformation of the histology image, and it indicates the magnitude of the largest local compression and largest local dilation of the image in relative units (1 = no compression).

Figure D.1 suggests an optimal range for a between 0.4 and 0.6, corresponding to slightly more conservative deformations at a = 0.6 based on the Jacobian ranges. All values of a, except for 0.2 led to diffeomorphic transformations with Jacobians > 0.

